# Irreversible inhibition of TRF2_TRFH_ recruiting functions: a strategy to induce telomeric replication stress in cancer cells

**DOI:** 10.1101/2023.02.01.526591

**Authors:** Alexander P. Sobinoff, Salvatore Di Maro, Ronnie R. J. Low, Rosaria Benedetti, Stefano Tomassi, Antonia D’Aniello, Rosita Russo, Ilaria Baglivo, Ugo Chianese, Paolo V. Pedone, Angela Chambery, Anthony J. Cesare, Lucia Altucci, Hilda A. Pickett, Sandro Cosconati

**Affiliations:** Telomere Length Regulation Unit, Children’s Medical Research Institute, Faculty of Medicine and Health, University of Sydney, Westmead, NSW 2145, Australia; DiSTABiF, University of Campania “Luigi Vanvitelli”, Via Vivaldi 43, 81100 Caserta, Italy; Genome Integrity Unit, Children’s Medical Research Institute, Faculty of Medicine and Health, University of Sydney, Westmead, NSW 2145, Australia; Department of Precision Medicine, University of Campania “Luigi Vanvitelli”, Vico L. De Crecchio 7, 80138, Naples, Italy; Department of Pharmacy, University of Naples “Federico II”, Via D. Montesano 49, 80131 Naples, Italy; BIOGEM, 83031 Ariano Irpino, Italy

## Abstract

The shelterin component telomeric repeat-binding factor 2 (TRF2) is an essential regulator of telomere homeostasis and genomic stability. Mutations in the TRF2_TRFH_ domain physically impair t-loop formation and prevent the recruitment of several factors that promote efficient telomere replication, resulting in a telomeric DNA damage response. Here, we design, synthesize, and biologically test covalent cyclic peptides that irreversibly target the TRF2_TRFH_ domain. We identify APOD53 as our most promising compound. APOD53 forms a covalent adduct with a reactive cysteine residue present in the TRF2_TRFH_ domain and induces phenotypes consistent with TRF2_TRFH_ domain mutants. These include induction of a telomeric DNA damage response in the absence of fusions, increased telomeric replication stress, and impaired recruitment of regulator of telomere elongation helicase 1 (RTEL1) and structure-specific endonuclease subunit (SLX4) to telomeres. We demonstrate that APOD53 impairs cell growth in both a telomerase-positive and an ALT cell line, while sparing the viability of non-cancerous cells. Finally, we find that co-treatment with APOD53 and the G4 stabilizer RHPS4 further exacerbates telomere replication stress.

## Introduction

Semiconservative DNA replication cannot replicate the ends of linear chromosomes entirely, resulting in the loss of DNA during each cell division (1). In addition, chromosome ends resemble DNA double-strand breaks that can potentially activate a DNA damage response (DDR) and cause genomic instability (2,3). These complications are overcome by telomeres, tandem arrays of TTAGGG nucleotide repeats terminating in a single-stranded G-rich 3’ overhang bound by the protective six-subunit complex shelterin (4). Telomeres buffer chromosome ends from genomic DNA erosion during semiconservative replication, while the shelterin complex inhibits the activation of an aberrant DDR.

Cumulative telomere erosion associated with cellular division eventually triggers a state of permanent cell cycle arrest known as cellular senescence (5,6). Bypass of senescence results in catastrophic telomere shortening, or telomere crisis (7). Shelterin function is lost in cells with critically short telomeres due to a lack of sufficient binding sites for this protective complex, resulting in widespread genomic instability (3,8). Cancer cells can escape from crisis by activating a telomere maintenance mechanism (TMM). Over 85% of cancers activate telomerase, whilst a smaller proportion activates the Alternative Lengthening of Telomeres (ALT) pathway (9,10). Despite their ability to maintain telomere length, cancer cell lines frequently display a DDR at their telomeres (11). Cancer cell lines that utilize the ALT pathway also exhibit characteristically high levels of telomeric replication stress (12). The aberrant telomeres of cancer cells, therefore, represent a vulnerability to be exploited for cancer therapy.

The shelterin component TRF2 is the primary suppressor of both the ataxia-telangiectasia mutated (ATM) kinase and non-homologous end joining (NHEJ) DNA damage response pathways at telomeres (2,13,14). TRF2 binds double-stranded telomeric DNA with high affinity and sequence specificity, acting as an essential recruiter of nucleases, helicases, and replication factors that function at telomeres. The TRFH domain of TRF2, which enables homodimerization of TRF2, is also essential for these recruiting functions (15). The TRF2_TRFH_ domain contains exposed lysine and arginine residues that allow dimeric TRF2 to wrap itself around telomeric DNA, thereby exerting topological stress (16). The TRF2_TRFH_ domain also promotes formation of the secondary telomere-loop (t-loop) structure that functionally conceals the chromosome end, preventing activation of ATM (17).

During S-phase, the telomere-associated helicase RTEL1 is recruited to telomeres by the TRF2_TRFH_ domain (18). RTEL1 is capable of unwinding telomeric G-quadruplexes with 5’–3’ polarity *in vitro* and is involved in t-loop disassembly during replication (19). The TRF2_TRFH_ domain also promotes efficient telomere replication by recruiting the endonuclease scaffold protein SLX4, whose association with the telomere peaks in late-S phase where it promotes the nucleolytic resolution of branched intermediates formed during replication (20). As TRF2 is responsible for the recruitment of several factors that promote efficient replication and prevent fork stalling, blocking the TRF2_TRFH_ domain has the potential to exacerbate the effects of replication stress-inducing treatments in cancer cells, thereby leading to mitotic catastrophe (21).

We have previously shown that targeting the TRF2_TRFH_ domain with a set of cyclic peptides induces a DDR in HeLa cancer cells (22). The best of these compounds (herein referred to as APOD41) displayed *in vitro* affinity for TRF2_TRFH_ in the sub-micromolar regimen (Kd = 0.12 μM) (22). Other groups have designed and synthesized novel peptides with demonstrated TRFH affinity *in vitro*, while no activity *in vivo* has been reported (23,24). In this study, we describe the structure-guided discovery of covalent cyclic peptides that irreversibly target the TRF2_TRFH_ domain by modifying our initial hit APOD41. We show that our most promising peptide, APOD53, is a proficient inducer of telomere dysfunction–induced foci (TIF), and inhibits both RTEL1 and SLX4 recruitment to the telomere. We further demonstrate that APOD53 forms a covalent adduct with the TRF2_TRFH_ domain and impairs cell growth in two cell models of telomere maintenance (telomerase-dependent and ALT), while sparing the viability of non-cancerous cells. When used in combination with the G4 stabilizer RHPS4 (25), APOD53 exacerbates telomere replication stress. This study reports the first covalent binder of the TRF2_TRFH_ domain, and the discovery of a new chemical genetics tool to study the role of TRF2-recruiting functions in the orchestration of telomere maintenance and telomere structural integrity.

## Methods

### Cell culture and cell lines

The cell lines HeLa, U-2 OS, and IMR-90 were were obtained from the American Type Culture Collection. HeLa and U-2 OS were cultured in Dulbecco’s modified Eagle’s medium (DMEM) supplemented with 10% (v/v) fetal bovine serum (FBS) in a humidified incubator at 37 °C with 10% CO2. IMR90 were cultured in DMEM supplemented with 1% (v/v) non-essential amino acids, 1% Glutamax (v/v), and 10% (v/v) FBS in a humidified incubator at 37°C with 10% CO2. The following comercial compounds were used in cell treatments: dimethyl sulfoxide (DMSO, Sigma-Aldrich), colcemid (Life Technologies), and Aphidicolin (Sigma-Aldrich). Cell lines were authenticated by 16-locus short-tandem-repeat profiling and tested for mycoplasma contamination by CellBank Australia (Children’s Medical Research Institute).

### Antibodies

A complete list of all primary antibodies (with manufacturers, catalog numbers and dilution factors) used throughout this study can be found in the appendix section (Appendix Table S1). Fluorophore-conjugated secondary antibodies (Life Technologies) were used for indirect immunofluorescence.

### Synthesis of RHPS4

3,11-Difluoro-6,8,13-trimethyl-8H-quino[4,3,2-kl]acridinium methyl sulphate (RHPS4) was synthesized as elsewhere reported (26). Briefly, 6-Fluoro-2-methylquinoline, 4 (2 g, 12 mmol) and dimethyl sulfate (1.11 ml, d: 1.333 g/ml, 12 mmol, 1 eq) were mixed and heated on a 100°C oil bath for 5 min. The crude product was recrystallized in methanol to give 1,2-Dimethyl-6-fluoro-quinolinium methyl sulfate, which was used in the next step without any further purification. Specifically, 1,2-Dimethyl-6-fluoro-quinolinium methyl sulfate (1.32 g, 4.5 mmol) and pyridine (0.46 ml, 4.5 mol, 1 eq) were dissolved in EtOH (60 ml) and the reaction was refluxed for 7 days. The crude product was crystallized from acetone to give 3,11-Difluoro-6,8,13-trimethyl-8H-quino[4,3,2-kl]acridinium methyl sulfate as a dark orange solid (0.12 g, 0.26 mmol, 11.5 % yield). The purity of the final compound was monitored by HLPC and its identity confirmed by HRMS (Figure s1).

### Synthesis of APOD41, APOD50-APOD55

Compounds APOD41, APOD50-APOD55 were synthesized on a Rink Amide AM-PS resin (182 mg, 0.1 mmol, 0.55 mmol/g) as solid support according to ultrasound-assisted Fmoc/*t*Bu solid-phase protocol (US-SPPS) previously reported by us (Figure s2)(27). Generally, Fmoc deprotections were carried out treating the solid phase with a 20% piperidine solution in DMF under ultrasound irradiation in a SONOREX RK 52 H cleaning bath (2 x 1 min.). Coupling reactions were performed employing 3 eq of amino acids preactivated with an equimolar amount of coupling reagents (COMU and OxymaPure) and 6 eq of DIPEA as a base (with respect to the resin functionalization) under ultrasound irradiation for 10 min.

Once on-resin linear elongation was completed, the Nα-Fmoc of the L-Tyr was removed and the free amino group was acetylated with acetic anhydride (38 μL, 4 eq) and DIPEA (140 μL, 4 eq) in DMF (4 ml) for 30 min. Next, the Allyl- and Alloc-protective groups of the L-Lys and D-Glu side chains were removed by treating the resin-bound peptides with a solution of 0.1 eq of tetrakis(triphenylphosphine)palladium(0) (12 mg) and 8 eq of DMBA (125 mg) in anhydrous DCM/DMF (2:1, 3 mL). The mixture was allowed to gently shake under argon atmosphere for 60 min and the treatment repeated once. The resin was drained, washed with DMF (3 times) and DCM (3 times), and treated with a solution of 0.06 M of potassium N,N-diethyldithiocarbamate in DMF (2 mL) for 60 min to remove any catalyst traces. The macrolactamization was performed using 3 eq of PyAOP and HOAt as dehydrating agents and 6 eq of DIPEA as a base in DMF for 6 hours.

Once the cyclization reaction was completed, APOD41 was cleaved from the solid support with a solution of TFA/TIS (95:5, 2 mL) for 3 h at room temperature. The exhausted resin was filtered, and the crude peptides precipitated from the cleavage solution diluting to 15 mL with cold Et_2_O, and then centrifuged (6000 rpm × 15 min). The supernatant was removed, and the crude precipitate resuspended in 15 mL of Et_2_O and then centrifuged once again. After removing the supernatant, the resulting solid was dried for 1 h under vacuum, dissolved in H_2_O/ACN (95:5), and purified by reverse-phase HPLC (solvent A: H_2_O + 0.1 % TFA; solvent B: ACN + 0.1 % TFA; from 10 to 70% of solvent B over 20 min, flow rate: 10 mL min-1). Fractions of interest were collected and evaporated from organic solvents, frozen, and then lyophilized. The obtained product was characterized by analytical HPLC (solvent A: H_2_O + 0.1 % TFA; solvent B: ACN + 0.1 % TFA; from 10 to 90% of solvent B over 20 min, flow rate: 1 mL min-1) and identity of peptides confirmed by high-resolution mass (Q Exactive Orbitrap LC-MS/MS (Thermo Fisher Scientific).

After the macrolactamization, the synthesis of APOD50-55 proceeded by reducing the azido group of the Pro residue with TCEP (86 mg, 0.3 mmol, 3 eq) in THF/H_2_O (9/1, v/v) for 3h at room temperature on an automated shaker. The resulting primary amine was then reacted with acryloyl chloride (25 μl, 0.3 mmol, 3 eq) in the presence of DIPEA (105 μl, 0.6 mmol, 6 eq) in DMF for 30 min (APOD50-APOD53) or chloroacetyl chloride (22 μl, 0.3 mmol, 3 eq) in the presence of DIPEA (105 μl, 0.6 mmol, 6 eq) in DMF for 30 min (APOD51-APOD54) or Cyanoacetic acid (103 mg, 0.36mmol, 3 eq) in the presence of COMU (103 mg, 0.36mmol, 3 eq) OxymaPure (103 mg, 0.36mmol, 3 eq) as coupling agents and DIPEA (105 μl, 0.6 mmol, 6 eq) in DMF for 60 min (APOD52-APOD55). Finally, the cyclopeptides were cleaved from the solid support, collected, purified, and characterized as above-described for APOD 41.

### Immnofluorescence and telomere-FISH

Indirect immunofluorescence and telomere-FISH were performed on both interphase nuclei and metaphase spreads. For metaphase spreads, cell cultures were treated with 20 ng/mL colcemid for 4 hrs, harvested by trypsinization, resuspended in 0.2% (w/v) KCl and 0.2% (w/v) trisodium citrate hypotonic buffer at 37°C for 5 min, and cytocentrifuged onto SuperFrost Plus glass slides (Menzel-Glaser) at 450g for 10 min in a Shandon Cytospin 4 at high acceleration. For interphase nuclei, cells were grown on coverslips. Cells on slides/coverslips were subjected to pre-extraction by incubation in KCM permeabilization solution (120 mM KCl, 20 mM NaCl, 10 mM Tris, 0.1% (v/v) Triton X-100) for 10 min. Cells were then washed in PBS, fixed at room temperature for 10 min in PBS with 4% (v/v) formaldehyde and blocked with 100 μg/mL DNase-free RNase A (Sigma) in antibody dilution buffer (20 mM Tris-HCl, pH 7.5, 2% (w/v) BSA, 0.2% (v/v) fish gelatin, 150 mM NaCl, 0.1% (v/v) Triton X-100 and 0.1% (w/v) sodium azide) for 1 hrs at room temperature. Cells were incubated with primary antibody diluted in antibody-dilution buffer for 1 hrs at room temperature, washed in Phosphate Buffered Saline Tween-20, and incubated with secondary antibody diluted in antibody dilution buffer for 1 hrs at room temperature. Coverslips were rinsed with PBS then fixed with 4% (v/v) formaldehyde at room temperature prior to telomere FISH. Coverslips were subjected to a graded ethanol series (75% for 2 min, 85% for 2 min, and 100% for 2 min) and allowed to air-dry. Dehydrated coverslips were overlaid with 0.3 μg/ml TAMRA–OO-(CCCTAA)_3_ or 0.3 μg/mL Alexa 488–OO-(CCCTAA)_3_ telomeric PNA probe (Panagene) in PNA hybridization solution (70% deionized formamide, 0.25% (v/v) NEN blocking reagent (PerkinElmer), 10 mM Tris–HCl, pH 7.5, 4 mM Na2HPO4, 0.5 mM citric acid, and 1.25 mM MgCl2), denatured at 80 °C for 5 min, and hybridized at room temperature overnight. Coverslips were washed twice with PNA wash A (70% formamide, 10 mM Tris pH 7.5) and then PNA wash B (50 mM Tris pH 7.5, 150 mM NaCI, 0.8% Tween-20) for 5 min each. DAPI was added at 50 ng/ml to the second PNA wash B. Finally, coverslips were rinsed briefly in deionized water, air dried and mounted in DABCO (2.3% 1,4 Diazabicyclo (2.2.2) octane, 90% glycerol, 50 mM Tris pH 8.0) or ProLong^™^ Gold Antifade Mountant (Thermofisher Scientific). Microscopy images were acquired on a Zeiss Axio Imager microscope with appropriate filter sets.

### EdU detection

Cells were pulsed with 10 μM EdU for 30 min. Cells were permeabilized, then fixed with 4% formaldehyde PBS solution. The Click-iT^®^ Alexa Fluor 647 azide reaction (Invitrogen) was then performed according to the manufacturer’s instructions.

### Genomic DNA extraction

Cells were harvested via trypsinization, washed in PBS, and resuspended in lysis buffer (50 mM Tris-HCl, 100 mM NaCl, 50 mM ethylenediamine tetraacetic acid (EDTA), 0.5% SDS, pH 8). Lysed cells were subjected to RNase A (50 μg/ml) treatment for 20 min at room temperature, followed by protein digestion with 400 μg/ml of proteinase K (Invitrogen) overnight at 55°C. DNA was extracted using two rounds of phenol/chloroform extraction followed by ethanol precipitation.

### C-circle assay

C-circles were amplified with Phi29 polymerase using dATP, dTTP, and dGTP overnight. Products were dot blotted onto Biodyne B membranes (Pall) and pre-hybridized in PerfectHyb Plus (Sigma) for at least 30 min. γ-[32P]-ATP-labeled telomeric C-probe (CCCTAA)4 was then added and blots were hybridized overnight at 37 °C (28). Blots were washed with 0.5× SSC, 0.1% SDS three times for 5 min each then exposed to a PhosphorImager screen. Imaging was performed on the Typhoon FLA 7000 system (GE Healthcare) with a PMT of 750 V.

### Immunoprecipitation (IP)

Cells were lysed in buffer A (20 mM HEPES-KOH pH 8, 0.3 M KCl, 2 mM MgCl2, 0.1% (v/v) Triton X-100, 10% (v/v) glycerol) supplemented with 1 mM PMSF (Cell Signaling) and 250 U/mL Benzonase (Novagen) for 2 h at 4 °C. Following lysis, the lysate was cleared at 16,000 × g for 30 min. Supernatant was then mixed with 10 μg of α-TRF2 (NOVUS) or 10ug α-IgG (Cell Signaling) and 20 μL Protein G/agarose beads overnight at 4 °C. The beads were then washed with 10 mL buffer A using vacuum suction and eluted with 1 × LDS loading buffer (Life Technologies) for 10 min at 70 °C.

### Western blot

Cells were collected and lysed in RIPA buffer (50 mM Tris-HCl pH 7.6, 150 mM NaCl, 1% Nonidet P-40, 0.5% sodium deoxycholate, 0.1% SDS, 4 mM EDTA) supplemented with cOmplete Mini EDTA-free protease inhibitor cocktail (Roche). Proteins were resolved on a 4%–12% bis-tris gel (Invitrogen). Transferred membranes were blocked in 5% milk and incubated with primary antibody overnight at 4 °C. Membranes were then incubated with corresponding HRP-conjugated secondary antibodies (Dako) for 1 h at room temperature, and bands visualized using PICO enhanced chemiluminescence reagents (Thermo Scientific).

### WST-1 cell viability assay

The WST-1 assay (Roche) was performed according to the manufacturer’s instructions. Briefly, cells were cultured in 96-well plates for up to 96 hrs. 10 μL of WST-1 reagent was added to each well and incubated for 30-60 min in standard culture conditions. The absorbance was then measured at OD = 420 nm.

### Protein expression and purification

BL21 DE3 *E. Coli* strain was transformed with TRF2_TRFH_-pet 15b plasmid containing the coding sequence for TRF2TRFH protein fragment to be expressed as His-tagged protein (22). Transformed colonies were inoculated in Luria-Bertani medium and grown at 37 °C until the optical density at 600 nm (OD600nm) reached 0.4. At this point of bacterial growth, 1 mM IPTG was added and the expression of the protein was carried out at 28 °C until the OD600nm reached 1.0. The His-tagged-TRF2_TRFH_ protein was purified in presence of cOmplete™ EDTA-free Protease Inhibitor Cocktail (Roche) by using Protino ^®^ Ni-TED 2000 Packed Columns (Macherey-Nagel) following the manufacturer’s protocol. The His-tag was removed by using Thrombin Protease (GE) at a concentration of 10 U/mg of protein. Thrombin cleavage was carried overnight at 4 °C. To purify the cleaved protein from the His-tag, a gel filtration chromatography was carried out by using a HiLoad 26/60 Superdex 75 gel filtration chromatography column (GE Healthcare, Milan, Italy) equilibrated with the following buffer: 25 mM Tris-HCl pH 8, 150 mM NaCl, 5mM DTT. The purified protein was checked by SDS-page after each step of the purification process.

### Mass spectrometry analysis

For MS analyses, preliminary experiments were carried out to determine the optimal protein concentration and APOD53 reagent-to-protein molar ratio to detect the covalent adduct. Under optimized conditions, aliquots of purified recombinant TRF2_TRFH_ (1 nmol in 50 μL of 25 mM Tris-HCl pH 8.0, 1 mM DTT) were incubated for 1h at room temperature in the presence of APOD53 with a 40-fold excess (mol/mol) and then subjected to enzymatic hydrolyses. The same amount of TRF2_TRFH_ without APOD53 was also digested as a control sample. Following reduction with 10 mM DTT (final concentration) for 1h at 56°C, an alkylation step was performed by incubating samples with 7.5 mM iodoacetamide (final concentration) at room temperature in the dark for 15. Tryptic and chymotryptic hydrolyses were then performed by adding bovine TPCK-treated trypsin (1 μg/μL, Sigma) or TLCK-treated chymotrypsin (1 μg/μL, Sigma) at a final enzyme:ligand ratio of 1:50 (w/w) and by incubating the mixtures overnight at 37 °C. Protein digestions were blocked by adding 1 μL of 2% trifluoroacetic acid (TFA) in water.

Following enzymatic digestions, samples were centrifuged (15800 g, 15 min). For matrix-assisted laser desorption ionization time-of-flight (MALDI-TOF) MS analyses, 1 μL of the peptide digests were mixed with 1 μL of saturated α-cyano-4-hydroxycinnamic acid matrix solution [10 mg/mL in acetonitrile/water (1:1, v/v), containing 0.1% TFA]. Thus, a droplet of the resulting mixtures (1 μL) was placed on the MALDI-TOF micro MX (Waters, Manchester, UK) target plate and dried at room temperature. Once the liquid was completely evaporated, samples were loaded into the mass spectrometer and analyzed. The instrument was externally calibrated by using a tryptic alcohol dehydrogenase digest (Waters) in reflectron mode. For linear mode, a four-point external calibration was applied using an appropriate mixture (10 pmol/μL) of ACTH fragment (18–39), insulin, cytochrome C, and horse myoglobin as standard proteins (Sigma). All spectra were processed and analyzed using MassLynx 4.1 software.

### Automated image analysis

ZEN microscopy images (.czi) were processed into extended projections of z-stacks using ZEN desk 2011 software (Zeiss) and imported into Cellprofiler v2.1.1 (29) for analysis. The DAPI channel was used to mask individual nuclei as primary objects. Foci within each segmented nucleus were identified using an intensity-threshold based mask. Any given object was considered to be overlapping another object when at least 80% of the first object’s area was enclosed within the area of a second object. For APBs, telomeres were considered to be overlapping with PML when at least 20% of the telomere was enclosed within PML.

### Statistical analysis

Statistical analysis was performed using GraphPad Prism. Box plots are displayed using the Tukey method where the box extends from the 25th to the 75th percentile data points and the line represents the median. The upper whisker represents data points ranging up to the 75th percentile+ (1.5 × the inner quartile range), or the largest value data point if no data points are outside this range. The lower whisker represents data points ranging down to the 25th percentile– (1.5 × the inner quartile range), or the smallest data point if no data points are outside this range. Data points outside these ranges are shown as individual points. Error bars, statistical methods and n, are described in figure legends.

## Results

### Structure-based design of TRF2_TRFH_ binders

To design new chemical entities that covalently bind the TRFH domain of TRF2 we utilized our previously published cyclic peptide APOD41 (22). This peptide was designed by analyzing the published X-ray coordinates of this domain complexed with the C-terminus of Apollo (ApoTBM, Apollo 496–532, PDB 3BUA) (30) and with a fragment of SLX4 (SLX4TBM, SLX4 1014–1028, PDB 4M7C) (31). APOD41 establishes hydrophobic contacts with the apolar residues lining the protein binding site (Figure 1A). Given the absence of polar residues lining the TRF2TRFH domain that could potentially be exploited as anchoring points in the search of more potent binders, we decided to turn APOD41 into a covalent binder of TRF2. This decision was stimulated by the realization that in our interaction model a TRF2 cysteine residue (Cys160) is lying close to the proline residue of the binding peptide. This latter amino acid in APOD41 was functionalized in position four of the pyrrolidine ring with three different electrophile groups, namely the classical acrylamide Michael acceptor that binds nucleophiles like cysteines (derivatives APOD50 and APOD53), the highly reactive chloroacetylamide group (derivatives APOD51 and APOD54), and the 2-cyanoacetamide which is widely used in medicinal chemistry (derivatives APOD52 and APOD55) (Figure 1B). These reactive species have reportedly low off-target effects, making them ideal warheads of choice for targeted covalent binders (32). Peptide synthesis was carried out according to the recently reported ultrasound-assisted Fmoc/tBu solid-phase synthesis approach (27), with details of the synthesis of APOD50-55 reported in the supporting information.

**Figure 1.**
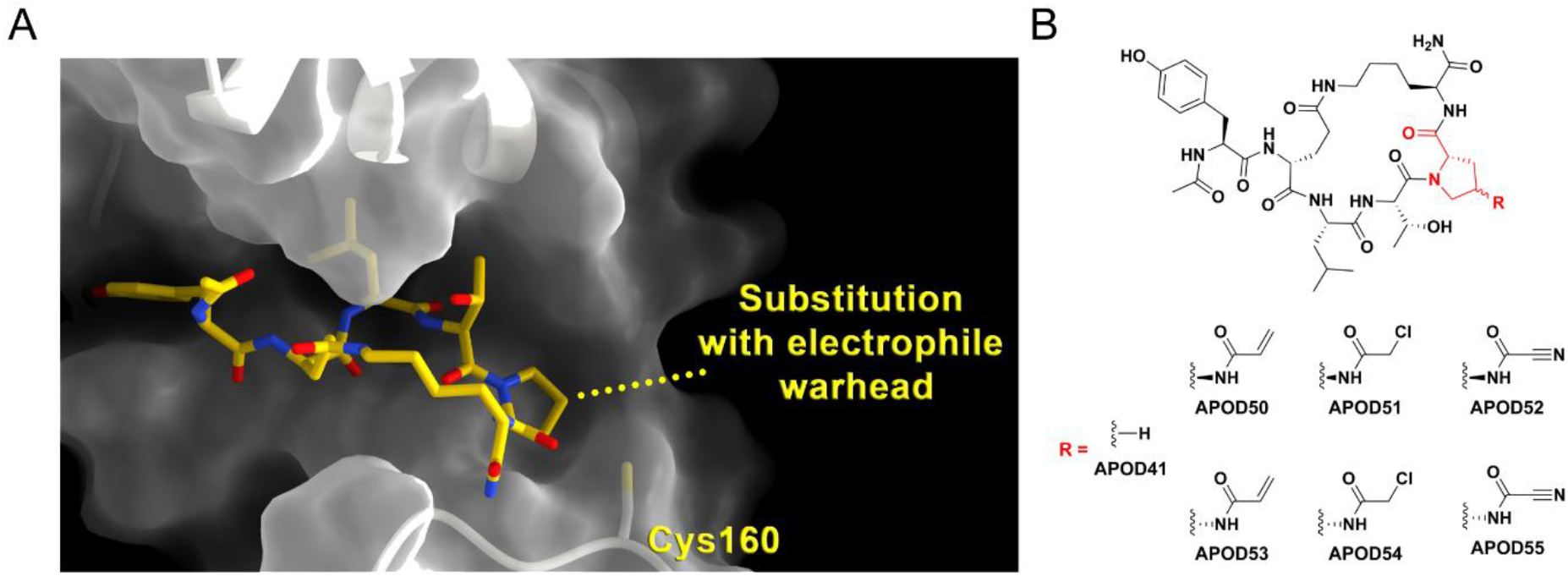
Structure-based design of TRF2_TRFH_ binders. (A) Binding pose of APOD41 into the three-dimensional structure of TRF2_TRFH_. The protein is represented as a transparent white surface and ribbons while the peptide as yellow sticks. The putatively reactive Cys residue is highlighted. (B) Chemical structures of APOD41 derivatives featuring the selected electrophile warheads.

### APOD41 derivatives induce telomeric dysfunction and alter ALT-associated phenotypes

To observe the effects of our covalent TRF2TRFH binding peptides on the DDR in cells, we exposed both HeLa (telomerase positive) and U-2 OS (ALT positive) cancer cells to the APOD41 derivative compounds (APOD50-55) and compared them to APOD41 and its stereoisomer STEREO8 (22), used here as a negative control. Following treatments, we quantified DDR-positive telomeres, termed “telomere deprotection-induced foci” or “TIF” (33) (Figure 2A). Consistent with previous findings, APOD41 induced TIF formation in both cell lines. Several of the APOD41 derivatives induced a greater telomere-specific DDR in both cell lines compared to APOD41, including APOD52, APOD53, and APOD54. The APOD41 derivatives also increased the total intensity of phosphorylated H2AX (γH2AX) in both HeLa and U-2 OS cells, indicative of a wider DDR (Figure s3A).

**Figure 2.**
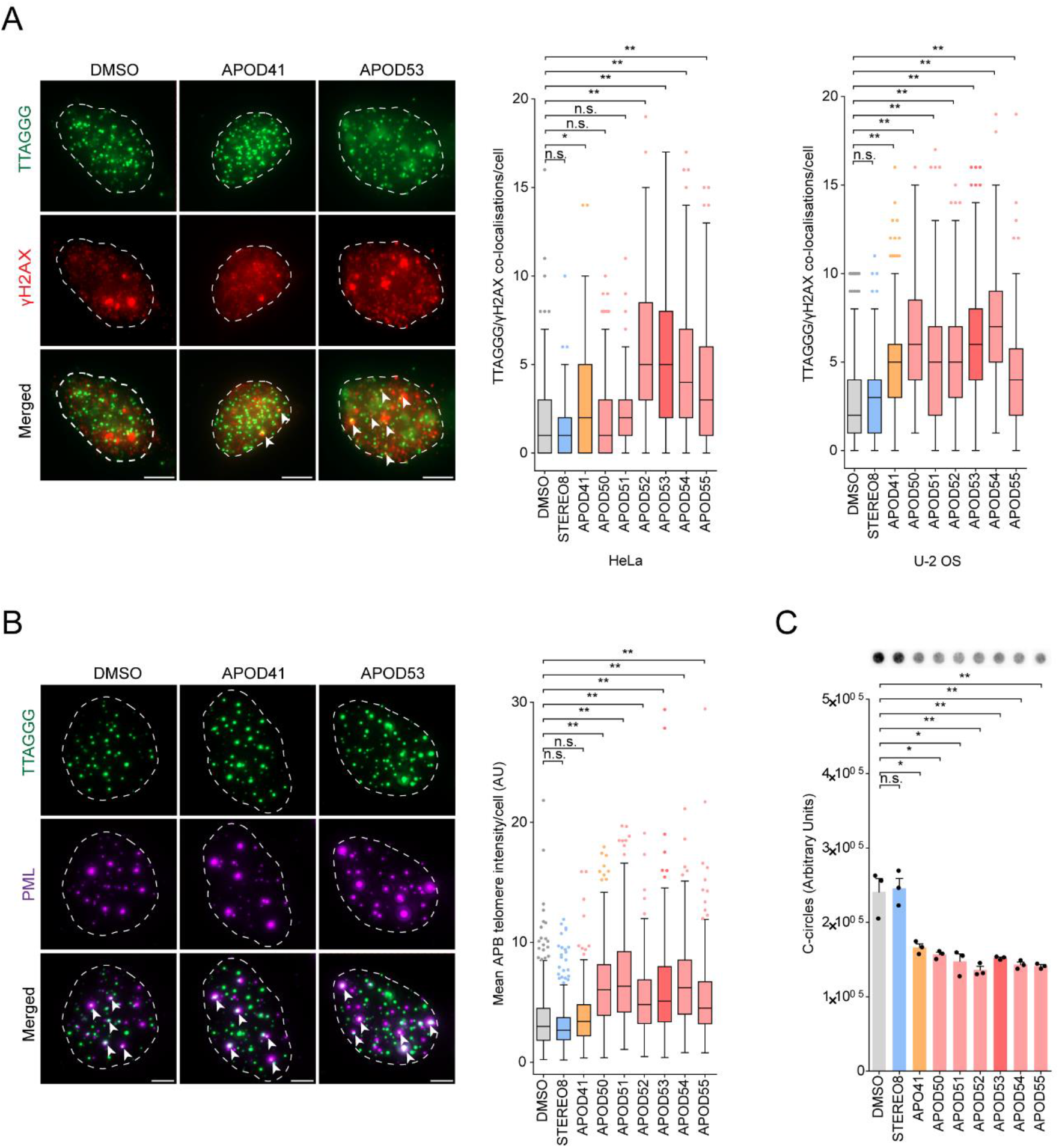
APOD41 derivatives induce a telomeric DDR and alter ALT associated phenotypes. (A) Representative images of telomere (green) and γ-H2AX (red) colocalizations (TIF) in HeLa cells treated with 1 μM of APOD41 derivatives for 24 hrs (left panel). TIF are indicated by white arrows. Scale bars are 5 μm. Tukey boxplots of TIF in HeLa and U-2 OS cells (right panel). Out of three experiments, *n* = 150 cells scored in HeLa and *n* = 220 cells scored in U-2 OS per treatment, n.s. = non-significant, \**p* < 0.05, \*\**p* < 0.01, Kruskal-Wallis test. (B) Representative images of telomere (green) and PML (purple) colocalizations (APBs) in U-2 OS cells treated with 1 μM of APOD41 derivatives for 24 hrs (left panel). APBs are indicated by white arrows. Scale bars are 5 μm. Tukey box plots of mean APB telomere foci intensity per cell. Out of three experiments, *n* ≥ 220 cells scored per treatment, n.s. = non-significant, \*\**p* < 0.005, Kruskal-Wallis test. (C) Representative dot blot and quantitation of C-circle assays in U-2 OS cells treated with 1 μM of APOD41 derivatives for 24 hrs. C-circle levels were normalized to the mean of DMSO control. Error bars represent the mean ± SEM from *n* = 3 experiments, n.s. = non-significant, \**p* < 0.05, \*\**p* < 0.01, Student’s *t*-test.

Cancers that utilize the ALT mechanism typically display elevated levels of telomeric replication stress and deprotection (11), which can be attributed to their dysregulated chromatin structure, variant telomeric sequences, and aberrant protein binding (34,35). Failure to alleviate this stress through fork regression and replication restart eventually triggers telomere synthesis by mechanisms analogous to break-induced replication (36). As the TRFH domain is responsible for recruiting factors that alleviate replication stress at the telomeres, we observed the effects of APOD41 derivatives on phenotypic markers of ALT activity. ALT-associated promyelocytic leukemia (PML) bodies (APBs) are a subset of PML bodies specific to ALT cells, in which telomeres aggregate and are potentially extended (37). Several APOD41 derivatives augmented the telomeric intensity within APBs, but did not drastically alter the number of APBs per cell, indicative of increased telomeric clustering (Figure 2B; Figure s3B).

We have observed this phenotype previously in response to increased replication stress at the telomere (38). APOD41 did not significantly alter these phenotypes.

ALT activity is also associated with the generation of extra-chromosomal telomeric repeat (ECTR) species, which can be both linear and circular in nature (39,40). The most frequently assayed circular species in ALT is partially single-stranded CCCTAA C-circles (28). Treatment with APOD41 and compounds APOD50-55 caused a significant reduction in C-circles (Figure 2C). The ability of APOD41 derivatives to repress C-circle production is intriguing, especially considering the paradoxical effect on APB telomeric intensity. However, the current lack of understanding regarding C-circle generation makes interpreting these results difficult.

### APOD53 impairs cancer cell growth

Of the APOD41 derivatives tested, APOD53 was chosen for further investigation due to the robustness and consistency in which it induced a DDR in both HeLa and U-2 OS cells. To determine the impact of APOD53 on cell viability, we examined the effects of gradated APOD53 exposure on cellular viability using the cell proliferation reagent WST-1 (41). APOD41 treatment caused an observable decrease in cell viability over time in both HeLa and U-2 OS cells, but only caused a statistically significant reduction in U-2 OS cells at 96 hrs (Figure 3). In contrast, treatment with its analog, APOD53, caused a greater and more significant decrease in cell viability at 48 hrs in HeLa and at 72 hrs in U-2 OS cells. Treatment with the APOD41 stereoisomer STEREO8, here used as negative control, did not affect cell viability in HeLa or U-2 OS cells.

**Figure 3.**
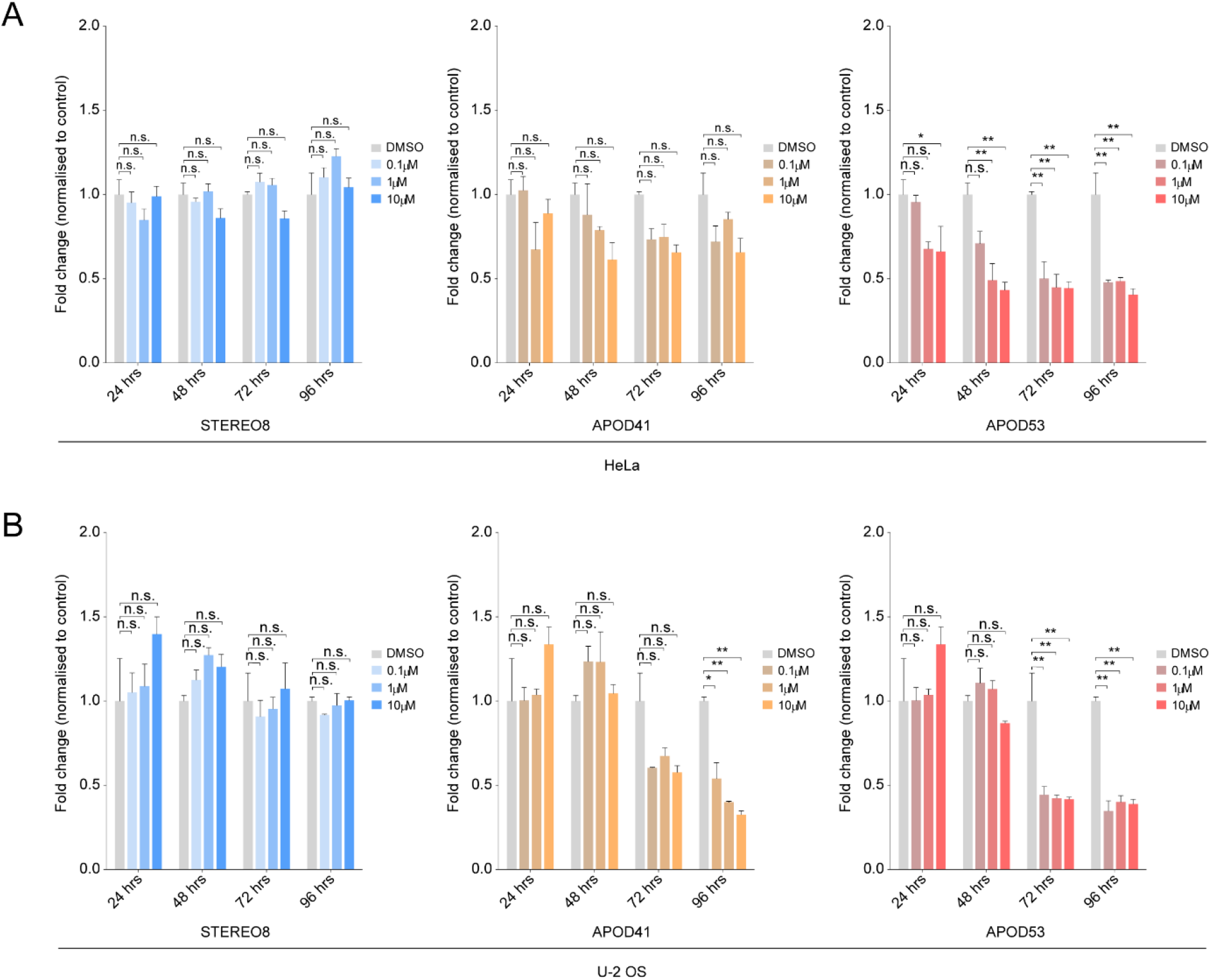
The APOD41 derivative APOD53 impairs cancer cell growth. Quantification of WST-1 cell proliferation assays in (A) HeLa and (B) U-2 OS with increasing dose over a 96 hr time period. Treated cells were normalized to the mean of DMSO control. Error bars represent the mean ± SEM from *n* = 3 experiments, n.s. = non-significant, \**p* < 0.05, \*\**p* < 0.01, ANOVA.

To further examine the effects of APOD53 on the telomeric DDR, we quantified metaphase TIF (mTIF) in untransformed IMR90 human fibroblasts (Figure s4A). Treatment with APOD53 caused a subtle increase in the DDR at telomeres, suggesting that exposure to these compounds preferentially affects cancer cell lines. WST-1 assays revealed that treatment with STEREO8, APOD41 and APOD53 all reduced IMR90 viability at early time points, but this reduction was not observed at longer treatment times (Figure s4B).

### APOD53 forms a covalent adduct with TRF2_TRFH_

To assess the covalent binding of APOD53, aliquots of recombinant TRF2_TRFH_ protein were analyzed by matrix-assisted laser desorption ionization time-of-flight (MALDI-TOF) MS following incubation with APOD53 and enzymatic proteolysis (Figure 4A). TRF2_TRFH_ without APOD53 was also analyzed as a control sample. MALDI-TOF profiles of TRF2_TRFH_ tryptic digestions were performed in both linear and reflectron mode to identify covalently modified TRF2_TRFH_ peptides. An ion signal at m/z 4326.19 was uniquely detected in the MALDI-TOF spectrum acquired in linear mode following incubation of TRF2_TRFH_ with APOD53 (Figure 4B). This ion corresponded to the covalent adduct (theoretical molecular mass, 4325.37 Da, [M+H]+=4326.38; Δ=0.19 Da) between TRF2_TRFH_ peptide 152-182 (152-IEEGENLDCSFDMEAELTPLESAINVLEMIK-182, average theoretical molecular mass, 3483.94 Da; numbering according to UniProtKB database, accession Q15554) and APOD53 (monoisotopic theoretical molecular mass, 841.43 Da). We next confirmed the presence of the covalent adduct by performing an additional MALDI-TOF MS analysis on chymotryptic digests (Figure 4C), revealing the presence of an ion signal at m/z 3341.29 in the spectrum of TRF2_TRFH_ incubated with APOD53. Indeed, this ion corresponded to the adduct (theoretical molecular mass, 3340.16 Da, [M+H]+= 3341.17; Δ=0.12 Da) between the TRF2_TRFH_ peptide 150-171 (150-SRIEEGENLDCSFDMEAELTPL-171, average theoretical molecular mass, 2498.73 Da) and APOD53. Overall, MALDI-TOF analyses confirmed the predicted site of covalent modification as Cys160.

**Figure 4.**
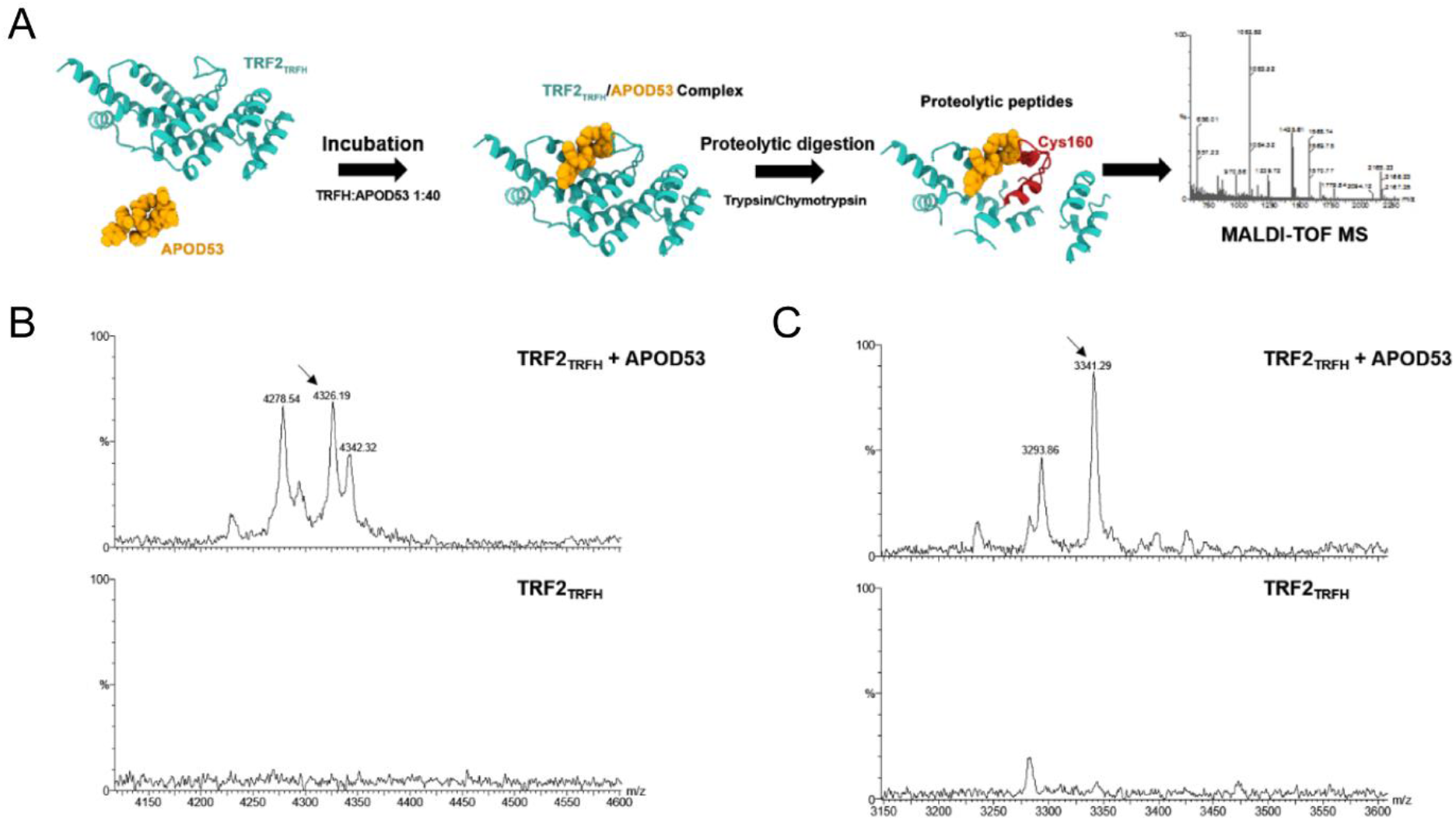
APOD53 forms a covalent adduct with TRF2_TRFH_. (A) Schematic workflow applied for investigating the covalent binding of APOD53 to TRF2_TRFH_. (B) Magnifications of MALDI-TOF spectra in the m/z range 4100-4600 of TRF2_TRFH_ tryptic peptides obtained in the presence (upper panel) and absence (lower panel) of APOD53. (C) Magnifications of MALDI-TOF spectra in the m/z range 3100-3600 of TRF2_TRFH_ chymotryptic peptides obtained in the presence (upper panel) and absence (lower panel) of APOD53. The covalent adducts of tryptic (ion signal at m/z 4326.19, sequence region 152-182) and chymotryptic (ion signal at m/z 3341.29, sequence region 150-171) peptides with APOD53 are indicated by arrows.

### Treatment with APOD53 induces telomere phenotypes associated with TRF2_TRFH_ inhibition

Mutants of TRF2 that lack the TRF2_TRFH_ domain or inhibit its function are unable to recruit the telomere-associated helicase RTEL1 (19) and the structure-specific endonuclease subunit SLX4 (42) to telomeres. To further validate APOD53 inhibition of the TRF2_TRFH_ domain we examined its effect on the ability of RTEL1 and SLX4 to interact with TRF2. The amount of RTEL1 and SLX4 co-immunoprecipitated with TRF2 was reduced in both HeLa and U-2 OS cells following treatment with APOD53 (Figure 5A). We also observed a significant decrease in the number of co-localizations between RTEL1 and telomeric DNA in both HeLa and U-2 OS cells following treatment with APOD53 (Figure 5B). In addition, the overall intensity of RTEL1 co-localizing with telomeres was significantly reduced (Figure 5C).

**Figure 5.**
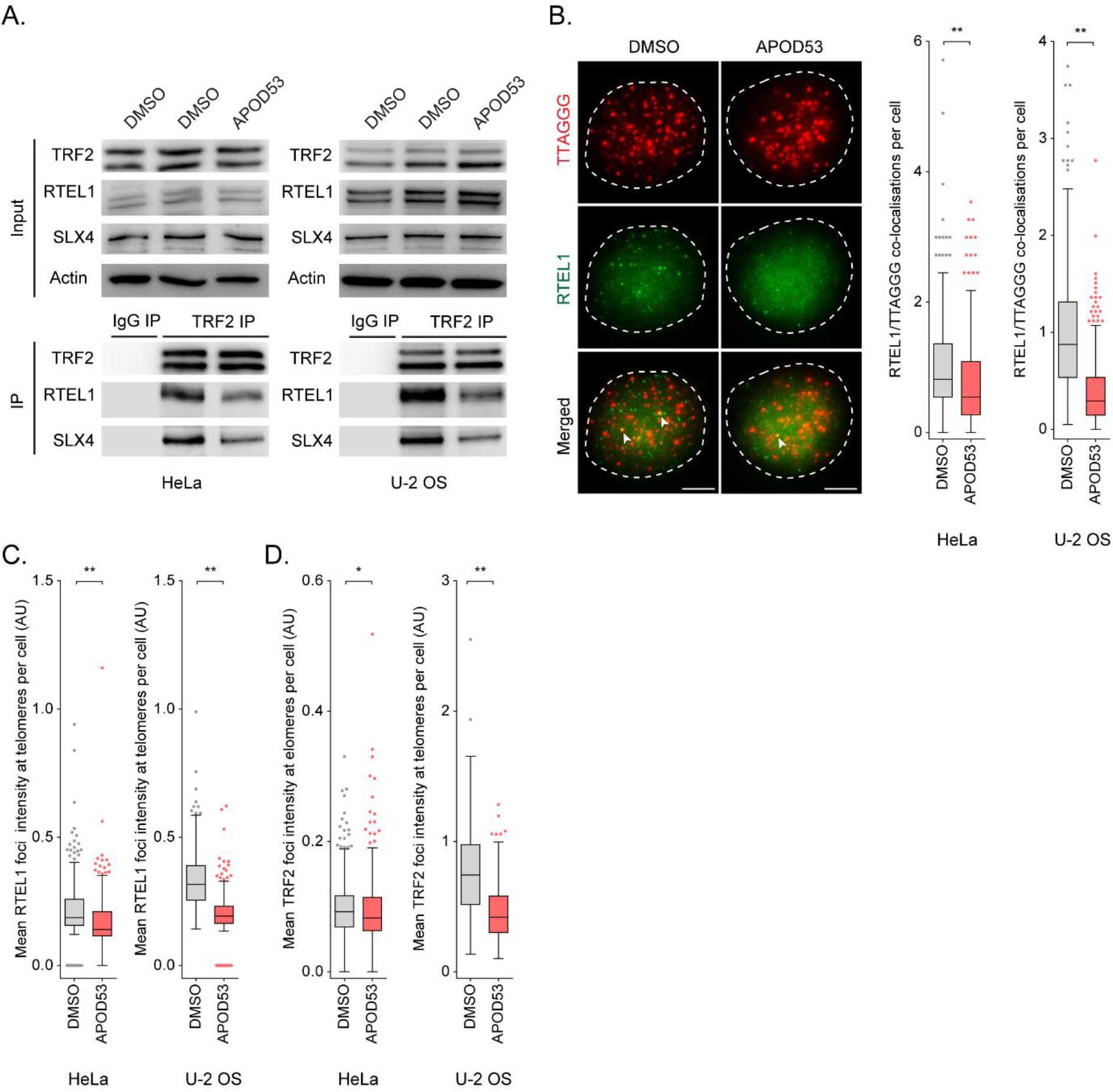
Treatment with APOD53 induces phenotypes consistent with TRF2_TRFH_ domain inhibition. (A) Interaction of TRF2_TRFH_ binding proteins, RTEL1 and SLX4, with endogenous TRF2 assayed by immunoprecipitation (IP) in HeLa and U-2 OS cells treated with 10 μM of APOD53 for 24 hrs. Input, indicated proteins assayed by western blot; IP, RTEL1 and SLX4 immunopurified using TRF2 antibodies assayed by western blot. IgG IP, non-specific IgG used for IP. (B) Representative images of telomere (red) and RTEL1 (green) co-localizations in U-2 OS cells treated with 10 μM of APOD53 for 24 hrs (left panel). RTEL1/TTAGGG co-localizations are indicated by white arrows. Scale bars are 5 μm. Tukey boxplots of RTEL1/TTAGGG co-localizations in HeLa and U-2 OS cells normalized to the average of DMSO (right panel). Out of three experiments, *n* = 414 cells scored in HeLa and *n* = 525 cells scored in U-2 OS per treatment, \*\**p* < 0.01, Mann–Whitney test. (C) Tukey boxplots of mean RTEL1 foci intensity at telomeres per cell in HeLa and U-2 OS cells treated with 10 μM of APOD53 for 24 hrs. Out of three experiments, *n* = 414 cells scored in HeLa and *n* = 525 cells scored in U-2 OS per treatment, \*\**p* < 0.01, Mann–Whitney test. (D) Tukey boxplots of mean TRF2 foci intensity at telomeres per cell in HeLa and U-2 OS cells treated with 10 μM of APOD53 for 24 hrs. Out of three experiments, *n* = 414 cells scored in HeLa and *n* = 525 cells scored in U-2 OS per treatment, n.s. = non-significant, \*\**p* < 0.01, Mann-Whitney test.

The TRF2^TRFH^ domain also promotes stable binding of TRF2 to the telomere by facilitating TRF2 homodimerization (15). Treatment with APOD53 did not affect TRF2 foci intensity at telomeres in HeLa cells, but did result in a significant reduction in foci in U-2 OS cells (Figure 5D). This suggests that TRF2 binding was compromised after APOD53 treatment in U-2 OS but not HeLa cells. As ALT cells have significantly longer telomeres on average compared to telomerase positive cells, combined with the abnormal nature of ALT cell telomeric chromatin, TRF2 binding may be more unstable at ALT telomeres, and therefore more susceptible to APOD53 inhibition. In accordance with other reports detailing TRF2_TRFH_ domain function (16,17,43), APOD53 did not induce NHEJ associated telomere fusions (data not shown).

### Co-treatment with APOD53 and a G4 stabilizer results in enhanced replication stress at telomeres

The TRF2^TRFH^ domain is an essential recruiter of several protein complexes involved in alleviating replication stress at telomeres. To determine whether APOD53 induces a defective replication phenotype at telomeres, cells were treated both independently and simultaneously with APOD53 and the G-quadruplex (G4) ligand, RHPS4, which has previously been reported to interfere with telomere replication (44). Analysis of HeLa and U-2 OS cells treated with APOD53 revealed enrichment of replication stress-induced pRPA2(S33) colocalizations at telomeres, with APOD53 inducing more pRPA2(S33)/TTAGGG colocalizations than low dose (0.4 μM) aphidicolin (APH) treatment in HeLa cells (Figure 6A). Unlike in U-2 OS, RHPS4 treatment alone did not induce pRPA2(S33) signaling at telomeres in HeLa cells. This could potentially be due to differences in telomere length/compaction between telomerase and ALT positive cells, with the longer de-compacted telomeres of ALT cells being more capable of forming G4 structures (45). Co-treatment with APOD53/RHPS4 caused a greater increase in the number of pRPA2(S33)/TTAGGG colocalizations per cell, compared to the individual treatments. Similarly, co-treatment with APOD53/APH increased the number of pRPA2(S33)/TTAGGG colocalizations per cell compared to cells treated with APH alone (Figure 6A). Although statistically significant, each treatment did not cause a drastic increase in the intensity of telomeric foci at pRPA2(S33)/TTAGGG co-localizations (Figure s5A). Treatment with APOD53 and RHPS4, both independently and simultaneously, increased the overall levels of pRPA2(S33) in HeLa and U-2 OS cells (Figure s5B), indicative of increased overall genomic replication stress.

**Figure 6.**
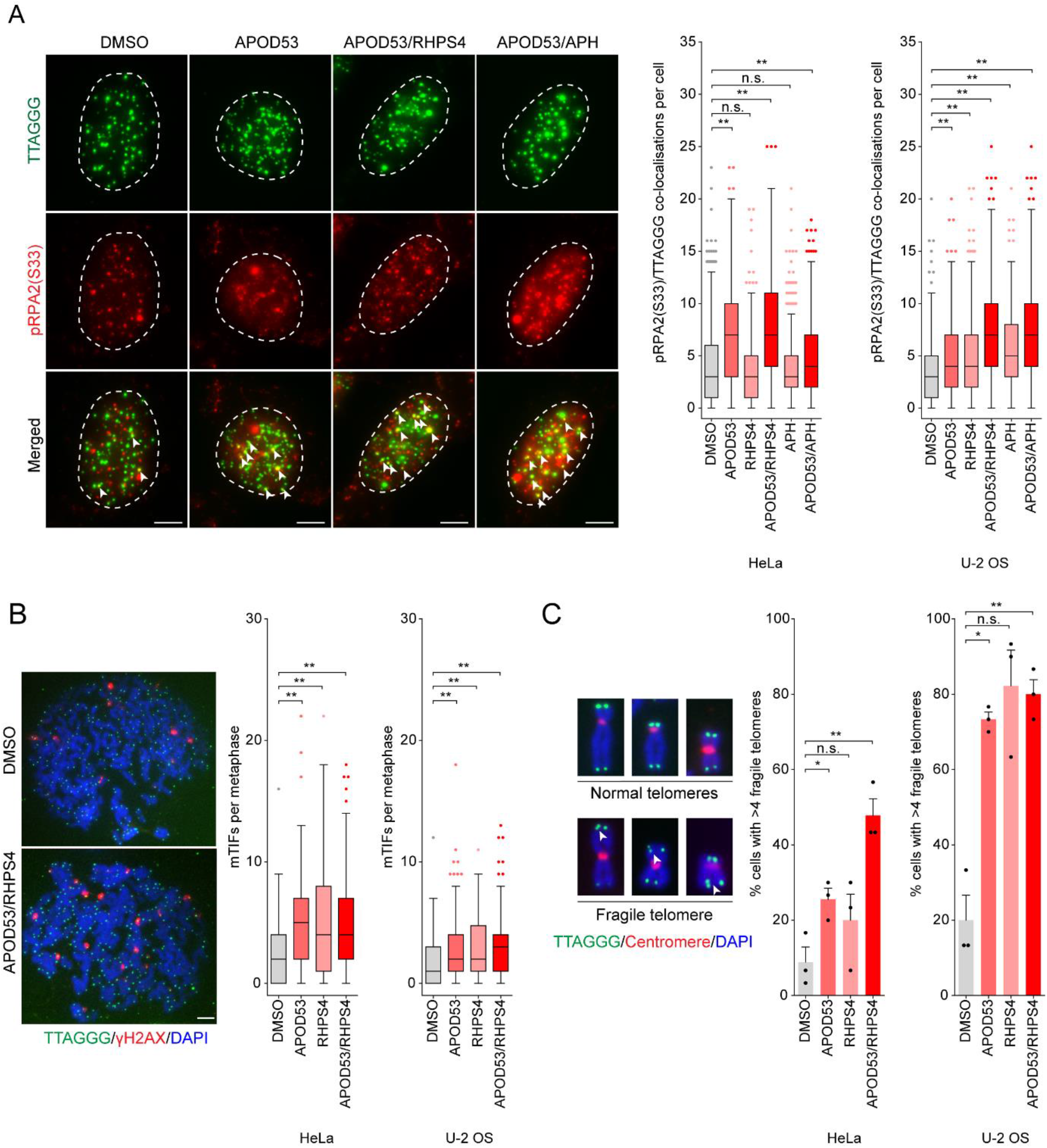
Co-treatment with APOD53 and a G4 stabilizer results in enhanced replication stress at the telomere. (A) Representative images of telomere (green) and pRPA2(S33) (red) colocalizations pRPA-TIF in U-2 OS cells treated with 10 μM of APOD53 and 1 μM RHPS4 for 24 hrs (left panel). pRPA-TIF are indicated by white arrows. Scale bars are 5 μm. Tukey boxplots of pRPA-TIF in HeLa and U-2 OS cells (right panel). Out of three experiments, *n* = 550 cells scored in HeLa and *n* = 600 cells scored in U-2 OS per treatment, n.s. = non-significant, \*\**p* < 0.01, Kruskal-Wallis test. (B) Representative images of telomere (green) and γ-H2AX (red) colocalizations on metaphase spreads (meta-TIF) in HeLa cells treated with 10 μM of APOD53 and 1 μM RHPS4 for 24 hrs (left panel). Scale bars are 5 μm. Tukey boxplots of meta-TIF. Tukey boxplots of meta-TIF in HeLa and U-2 OS cells (right panel). Out of three experiments, *n =* 120 metaphases scored per treatment. \*\**p* < 0.01, Kruskal-Wallis test. (C) Representative images of normal and fragile telomeres (double telomere signal) (left panel). Fragile telomeres are indicated by white arrows. Quantitation of fragile telomeres in HeLa and U-2 OS cells treated with 10 μM of APOD53 and 1 μM RHPS4 for 24 hrs (right panel). Error bars represent the mean ± SEM from *n* = 3 experiments, *n* = 120 metaphases scored per treatment, n.s. = non-significant, \**p* < 0.05, \*\**p* < 0.01, Student’s *t*-test.

53BP1 is a marker of DNA double strand breaks and NHEJ signaling. Treatment with either APOD53 or RHPS4 alone did not cause a significant increase in 53BP1/TTAGGG foci in HeLa cells (Figure s5C); however, co-treatment with APOD53/RHPS4 did increase 53BP1 signaling, suggesting a potential increase in ATM signaling. APOD53 and RHPS4 treatment alone or in combination did not affect 53BP1 signaling at U-2 OS telomeres, and did not induce telomere fusions in either HeLa or U-2 OS cells (data not shown).

We also observed an increase in mTIF consistent with heightened telomeric damage after APOD53, RHPS4 and APOD53/RHPS4 treatments in both HeLa and U-2 OS cells, with APOD53/RHPS4 co-treatment in U-2 OS cells producing the slightly greater effect on telomeric dysfunction (Figure 6B). As mTIF monitoring avoids the scoring of TIF postulated to transiently occur during the S/G2 phases, mTIF represent a more accurate estimate of telomeric damage in a whole nucleus (11,46). We next quantified the formation of fragile telomeres, multiple and/or highly extended telomeric fluorescence *in situ* hybridization (FISH) signals at individual chromatid ends detected in metaphase. Telomeres resemble common fragile sites, and fragile telomeres are quantified as specific markers of telomeric replication stress (47,48). APOD53, RHPS4 and APOD53/RHPS4 treatments all increased fragile telomeres in both HeLa and U-2 OS cells, with APOD53/RHPS4 co-treatment in HeLa cells causing the greatest induction in telomere fragility (Figure 6C).

To determine whether APOD53 exposure influences overall replication rates we quantitated EdU incorporation as a measure of genomic replication stress. In U-2 OS cells, we identified a higher number of EdU positive cells in both APOD53 and APOD53/RHPS4 treated groups compared to DMSO control, suggesting a higher proportion of cells in S-phase (Figure s6A). APOD53, RHPS4, and APOD53/RHPS4 treatments did not significantly decrease the rate of DNA replication as measured by EdU intensity in U-2 OS cells, as was observed following treatment with APH, suggesting the higher proportion of actively replicating cells was not due to S-phase stalling (Figure s6B). APOD53, RHPS4, and APOD53/RHPS4 treatment did not influence the number of EdU positive HeLa cells, although APOD53/RHPS4 treatment did significantly reduce the rate of EdU incorporation (Figure s6A; s6B). As expected, APH and APOD53/APH caused a significant increase in EdU positive cells and significantly decreased replication rates in both cell lines (Figure s6A; s6B).

As another marker of genomic replication stress/stability, we also assayed for micronuclei, small nuclei that form due to chromosome segregation errors during mitosis (49). Treatment with APOD53, RHPS4, and APOD53/RHPS4 did not significantly increase micronuclei formation, although the APOD53/RHPS4 treatment did cause an observable increase in micronuclei in HeLa cells (Figure s6C). As expected, treatment with a low dose of APH caused a significant increase in micronuclei formation.

Finally, cell viability assays were used to examine the combination effect of APOD53 and RHPS4 on cell viability. APOD53 had a more consistent effect on cell viability between HeLa and U-2 OS cells compared to RHPS4 at the chosen concentrations (Figure s7). The combination treatment did not produce an additive effect on viability, compared to APOD53 treatment alone, in either cell line.

## Discussion

The unlimited proliferative capacity of cancer cells is entirely dependent upon telomere maintenance, making factors that control telomere elongation attractive chemotherapeutic targets to limit cancer cell proliferative capacity. However, inhibition of telomere elongation has limitations, the most critical of which is the extended lag phase between treatment and effect as telomeres shorten to induce a second “crisis”. The telomere binding protein TRF2 is the principal shelterin component involved in telomere capping and the recruitment of factors that both promote efficient replication and prevent fork stalling (3). An alternative strategy to telomere maintenance inhibition could involve targeting TRF2 function in cancer. Novel compounds with the ability to down-regulate or inhibit TRF2 need to be developed to test this hypothesis.

We have previously identified the first cell-permeable TRF2 binding chemotype (APOD41) capable of targeting the TRF2_TRFH_ domain and inducing a DDR in cancer cell lines (22). To improve the effectiveness of our original compound, here we screened several APOD41 derivatives modified with reactive species to induce covalent binding. Turning this peptide into a covalent binder of TRF2 offers the opportunity to improve its biochemical efficiency by exploiting non-equilibrium binding. Moreover, irreversible interruption of the TRF2^TRFH^-recruiting functions by covalent binding should be more difficult to overcome by feedback mechanisms, such as overexpression of TRF2 or one of its interactors. Indeed, the majority of the APOD41 derivatives performed better than the parent compound by inducing a stronger telomeric DDR in cancer cell lines independent of the telomere maintenance mechanism, indicative of the modified covalent binders targeting the TRF2_TRFH_ domain more efficiently.

As the ALT mechanism can be impacted by several factors recruited to the telomere by TRF2 through the TRFH domain, such as RTEL1 (50) and SLX4 (42), we examined the effects of the APOD41 derivatives on the ALT phenotypes APBs and C-circles. Unlike the parent compound, the APOD41 derivatives increased telomere clustering in APBs. We have observed increased telomere clustering in APBs previously by inducing replication fork stalling through FANCM depletion (38), suggesting this phenotype may be due to replication stress. Interestingly, all the TRF2_TRFH_ binders caused a reduction in the production of C-circles. APB frequency and C-circle levels are often positively associated with each other, but a link between telomeric APB clustering and C-circles has not been reported. As the observed changes in C-circle levels were not drastic (< 2-fold) and the origin of the ECTRs is unknown, it is hard to speculate the biological significance of this result.

Further analysis revealed the cysteine binding acrylamide Michael acceptor modifier APOD53 can form covalent adducts with the TRF2_TRFH_ peptide by exploiting the free cysteine residue (Cys160) lying close to the hypothesized binding site on TRF2. We found APOD53 to be a more potent inhibitor of cellular viability than APOD41 in cancer cells, and that neither peptide caused long-term loss of viability of mortal IMR90 cells. This may be due to the increased replication stress inherent in all cancer cells making them more susceptible to TRF2 inhibition (51). Treatment with APOD53 recapitulated the effects of TRF2_TRFH_ defective mutants by blocking the interaction of RTEL1 and SLX4 with TRF2, and by inducing replication stress, DNA damage, and telomere fragility (18). Treatment with APOD53 was, however, not sufficient to induce telomere fusions that have previously been associated with the loss of TRF2. This is most likely due to incomplete suppression of TRF2 activity with APOD53, compared to almost complete deletion of TRF2 being required to induce the telomeric fusion phenotype (46), and that mutation of the TRFH domain inhibits t-loop formation but does not reduce the ability of TRF2 to inhibit NHEJ (16,17).

We further determined that co-treatment with the G4 stabilizing agent RHPS4 exacerbated the telomeric replication stress caused by APOD53 in both HeLa and U-2 OS cells. The levels of replication stress observed at the telomeres with combination treatments exceeded that of low dose APH treatments, whilst causing less overall genomic replication stress. This result can be attributed to APOD53 inhibiting the telomeric recruitment of multiple proteins involved in dismantling the G4 structures stabilized by RHPS4, including RTEL1 (18). We found both APOD53 and RHPS4 increased telomeric DNA damage and reduced cell viability, but, unlike the effects on the replication stress response, combined treatments did not exacerbate the effects. Indeed, RHPS4 treatment did not influence HeLa cell viability. These results suggest that changes in telomeric replication stress markers do not always directly translate into a parallel telomeric DDR. Indeed, we observed a higher baseline of fragile telomeres in DMSO treated U-2 OS cells compared to HeLa, and a higher baseline of mTIF in HeLa compared to U-2 OS. The lack of an additive effect could also suggest that G4 formation and TRF2^TRFH^ inhibition cause DNA damage signaling and reduce cell viability via the same pathway in U-2 OS/ALT cells, and that one of these two insults induces a response upstream of the other, but not in the case of HeLa/telomerase positive cell viability. This may be due to the inherent differences in telomere biology between ALT and telomerase positive cells, with U-2 OS/ALT cells being more susceptible to G4 formation due to their longer de-compacted telomeres and G4 induced replication stress due to their elevated levels of replication stress at baseline.

In conclusion, we have generated the first cell-permeable chemical genetics tool that covalently binds the TRF2_TRFH_ domain and accurately recapitulates inhibition of the TRF2_TRFH_ domain recruiting functions, demonstrating for the first time that covalent inhibition of the domain prevents recruitment of RTEL1 and SLX4 to TRF2 in cells. Our covalent binder APOD53 induces a stronger telomeric DDR and has a more pronounced inhibitory effect on cancer cell viability compared to our previous non-covalent compound APOD41. We also demonstrate that cancer cell lines may be more susceptible to covalent TRF2_TRFH_ domain inhibition than mortal cells, and that the telomeric replication stress induced by this inhibition can be exacerbated by stabilizing G4 structures in U-2 OS/ALT cells. These results provide the rationale for TRF2_TRFH_ domain inhibition as a potential cancer therapeutic target.

## Data availability statement

The authors declare that all data related to findings of this study are available within the article and its Supplementary Information files, or from the corresponding authors upon request.

