## Supplementary material for "Irreversible inhibition of TRF2_TRFH_ recruiting functions: a strategy to induce telomeric replication stress in cancer cells": Supllemental tables and figures

**Appendix Tables**

**Appendix Table S1. List of antibodies used in this study**

| Target | Species | Source, Cat # | Immunofluorescence Dilution | Western Blot Dilution |
| --- | --- | --- | --- | --- |
| γ-H2AX | Mouse | Millipore, 05-636 | 1.500 | - |
| PML | Goat | Santa Cruz Biotech, sc-9862 | 1.500 | - |
| RTEL1 | Rabbit | NOVUS, NBP2-22360 | 1.500 | 1.2000 |
| TRF2 | Mouse | Millipore, 05-521 | 1.500 | - |
| pRPA2(S33) | Rabbit | Bethyl Laboratories, A300-246A | 1.500 | - |
| 53BP1 | Rabbit | Santa Cruz Biotech, sc-22760 | 1.500 | - |
| TRF2 | Rabbit | NOVUS, NB110-57130 | - | 1.2000 |
| SLX4 | Rabbit | NOVUS, NBP1-28679 | - | 1.2000 |
| Actin | Rabbit | SIGMA, A2066 | - | 1.5000 |

**Supplementary figures**

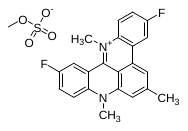
**RHPS4** 120 mg, yield: 11.5 %, purity: ≥ 95%, tR 18.34 min, (analytical HPLC, 30 to 90% acetonitrile (0.1% TFA) in water (0.1% TFA) over 10 min, flow rate of 1.0 mL/min); ESI-MS: Calculated: 347,14 for C_23_H_20_F_2_N_2_O_4_S [M]^+^, found: 347.05.

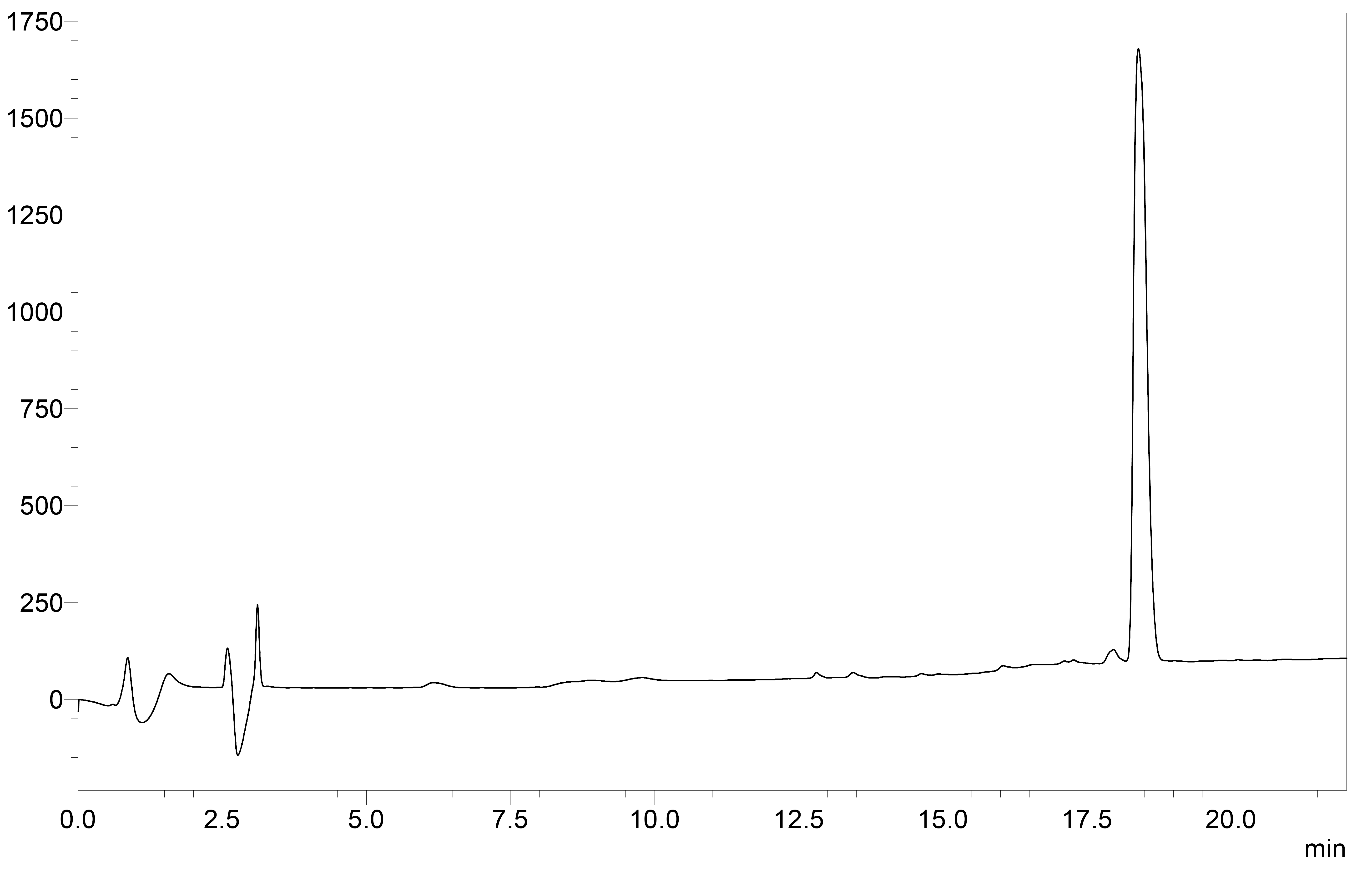

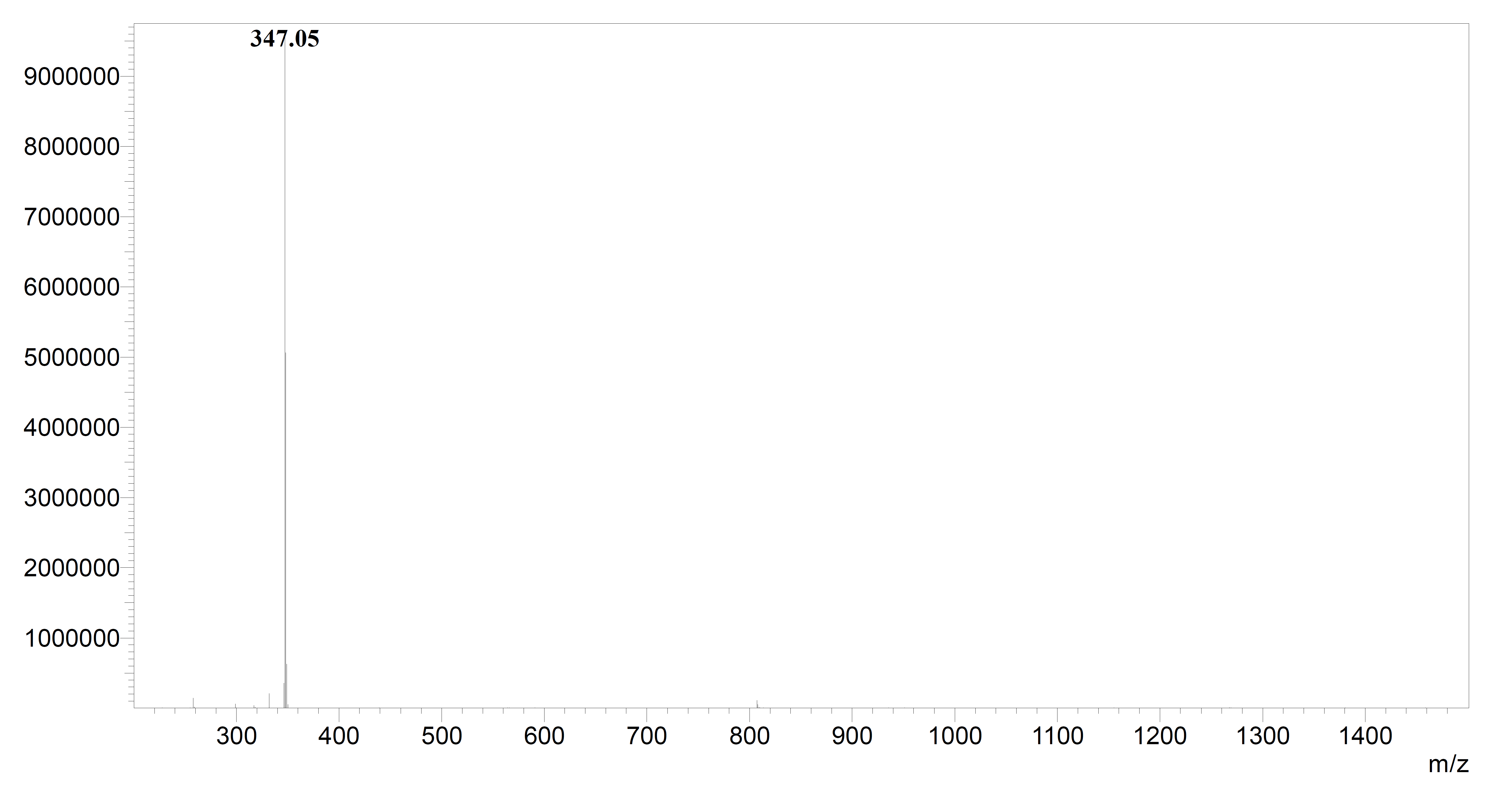

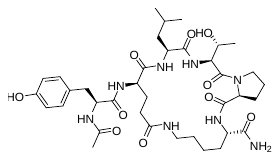
**APOD41** 43 mg, crude yield: 56 %, purity: ≥ 95%, tR 12.54 min, (analytical HPLC, 10 to 90% acetonitrile (0.1% TFA) in water (0.1% TFA) over 20 min, flow rate of 1.0 mL/min); ESI-MS: Calculated: 773.72 for C_37_H_57_N_8_O_10_ [M+H]^+^, found: 773.25.

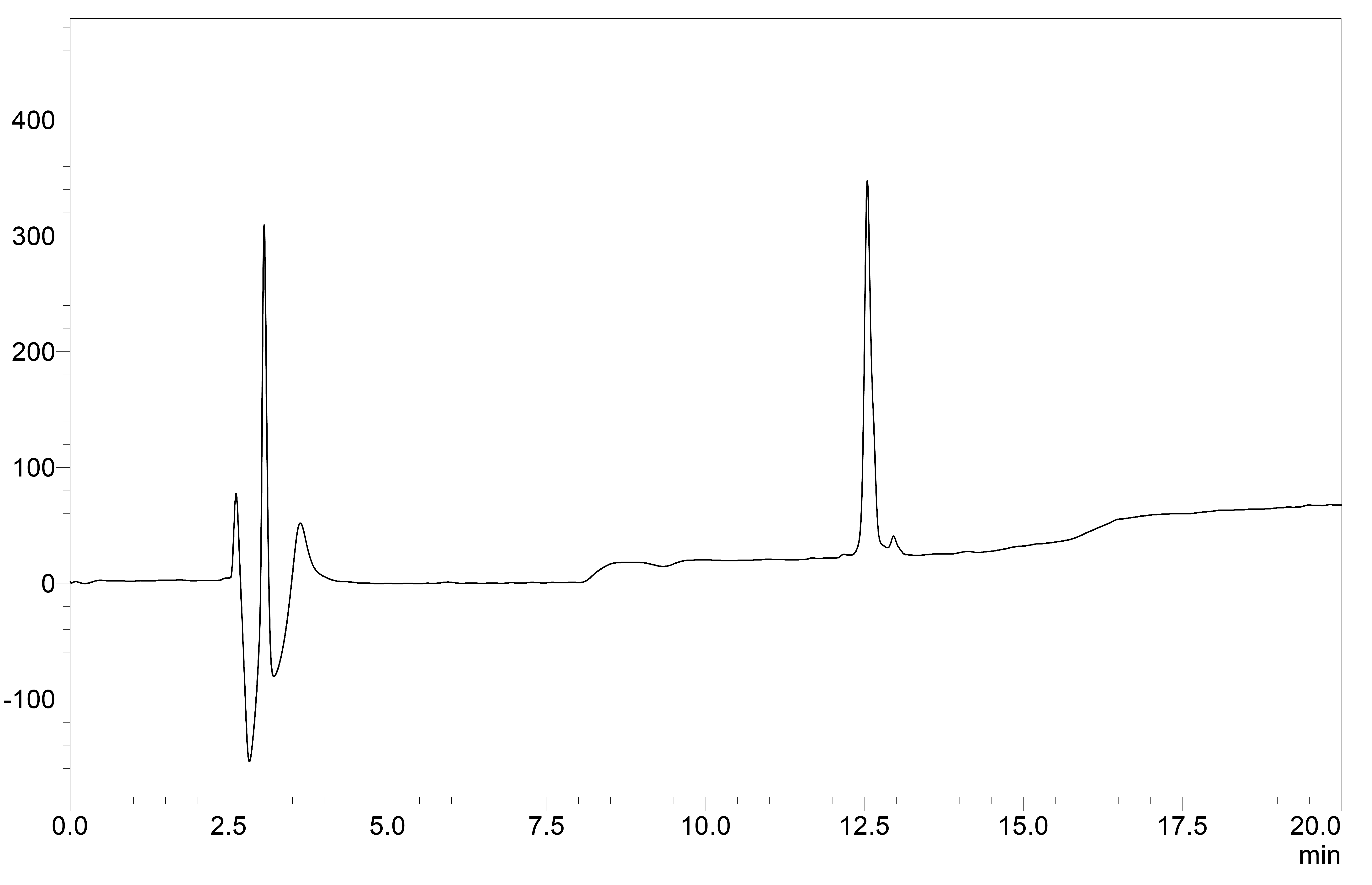

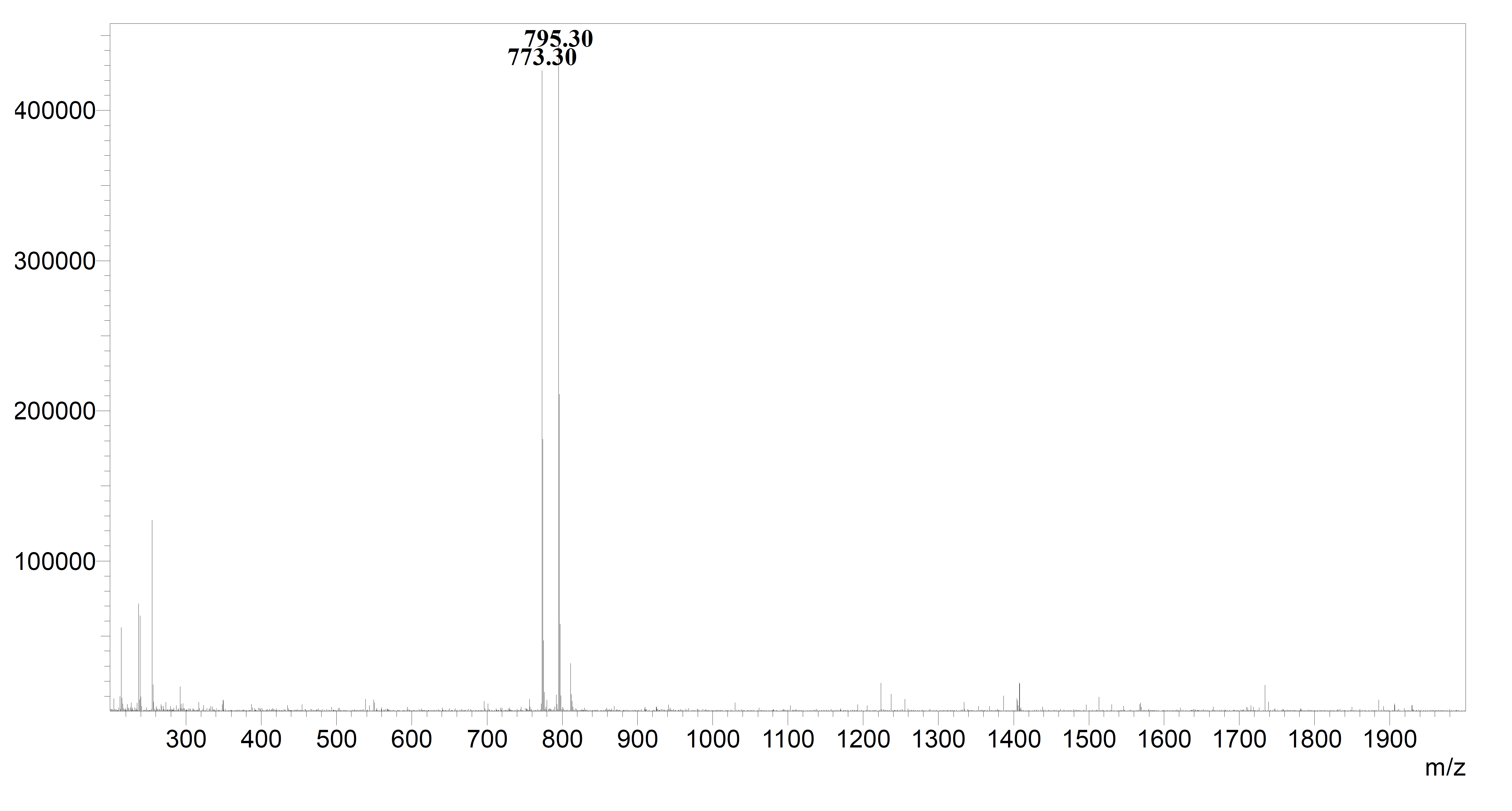

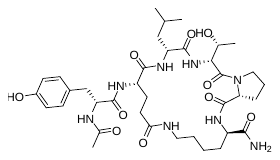
**STEREO8** 52 mg, crude yield: 67 %, purity: ≥ 95%, tR 12.59 min, (analytical HPLC, 10 to 90% acetonitrile (0.1% TFA) in water (0.1% TFA) over 20 min, flow rate of 1.0 mL/min); ESI-MS: Calculated: 773.72 for C_37_H_57_N_8_O_10_ [M+H]^+^, found: 773.40.

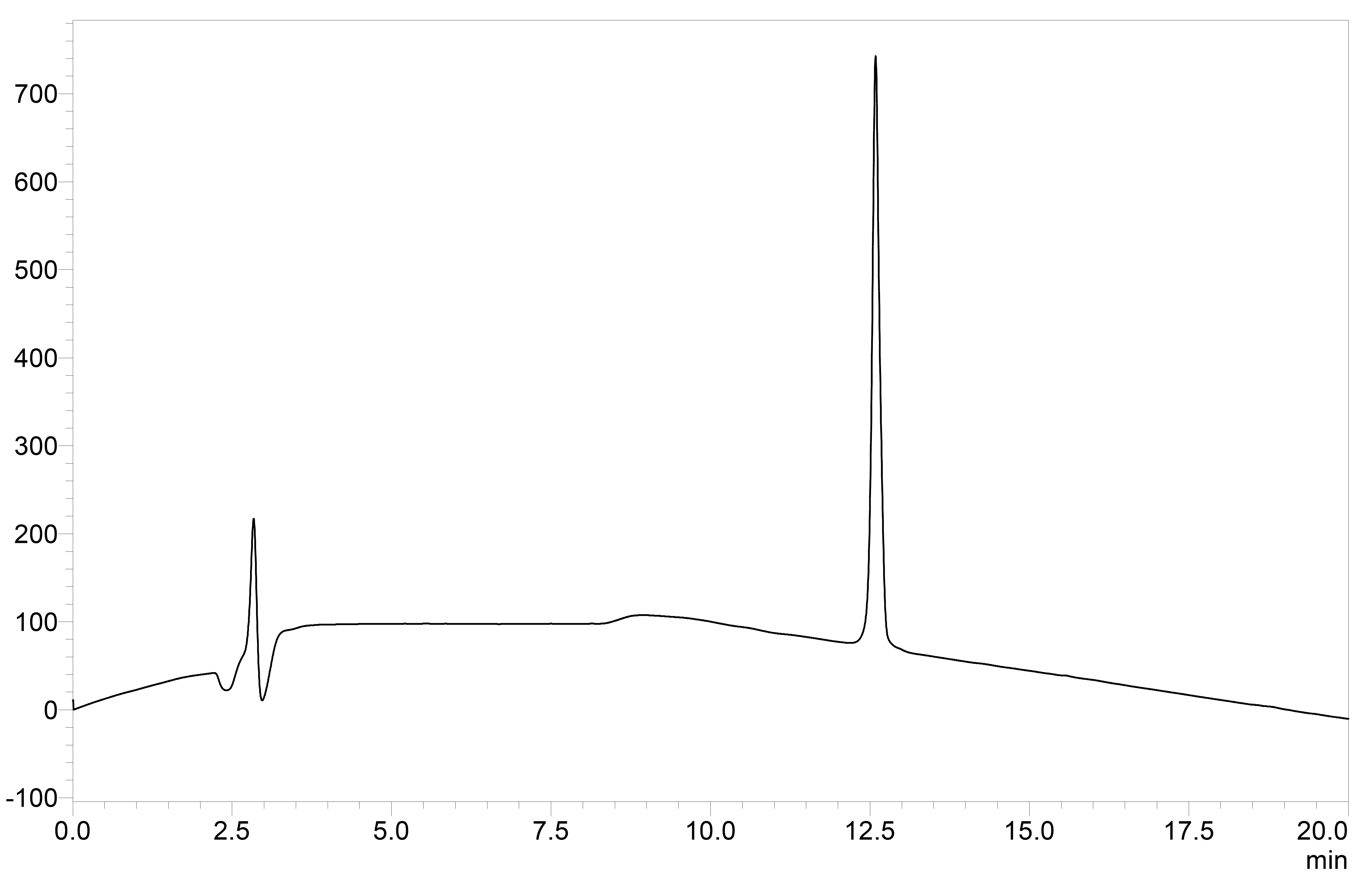

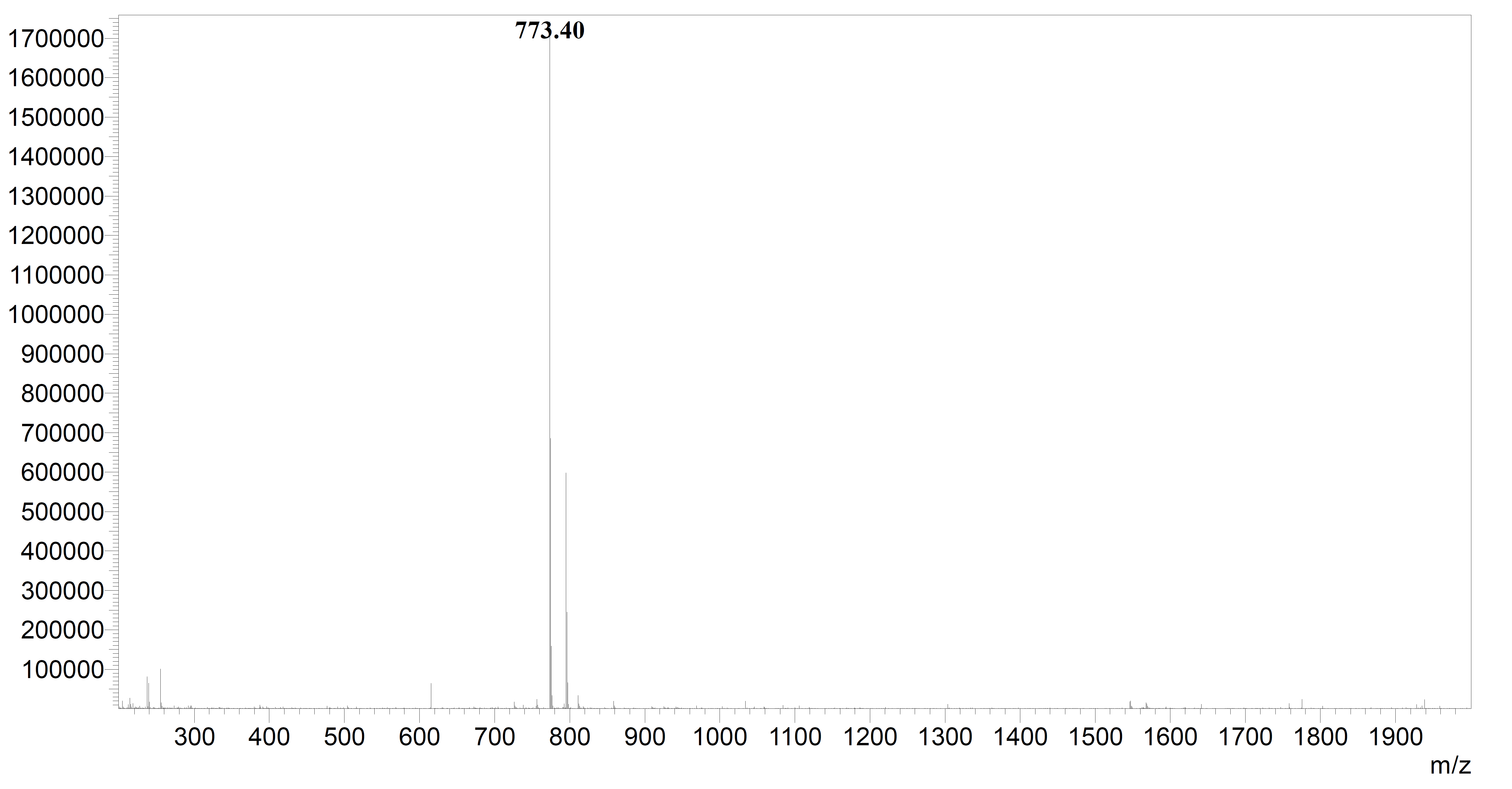

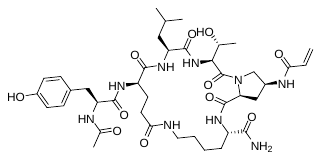
**APOD50** 34 mg, crude yield: 40 %, purity: ≥ 95%, tR 12.61 min, (analytical HPLC, 10 to 90% acetonitrile (0.1% TFA) in water (0.1% TFA) over 20 min, flow rate of 1.0 mL/min); ESI-MS: Calculated: 842.96 for C_40_H_60_N_9_O_11_ [M+H]^+^, found: 842.45.

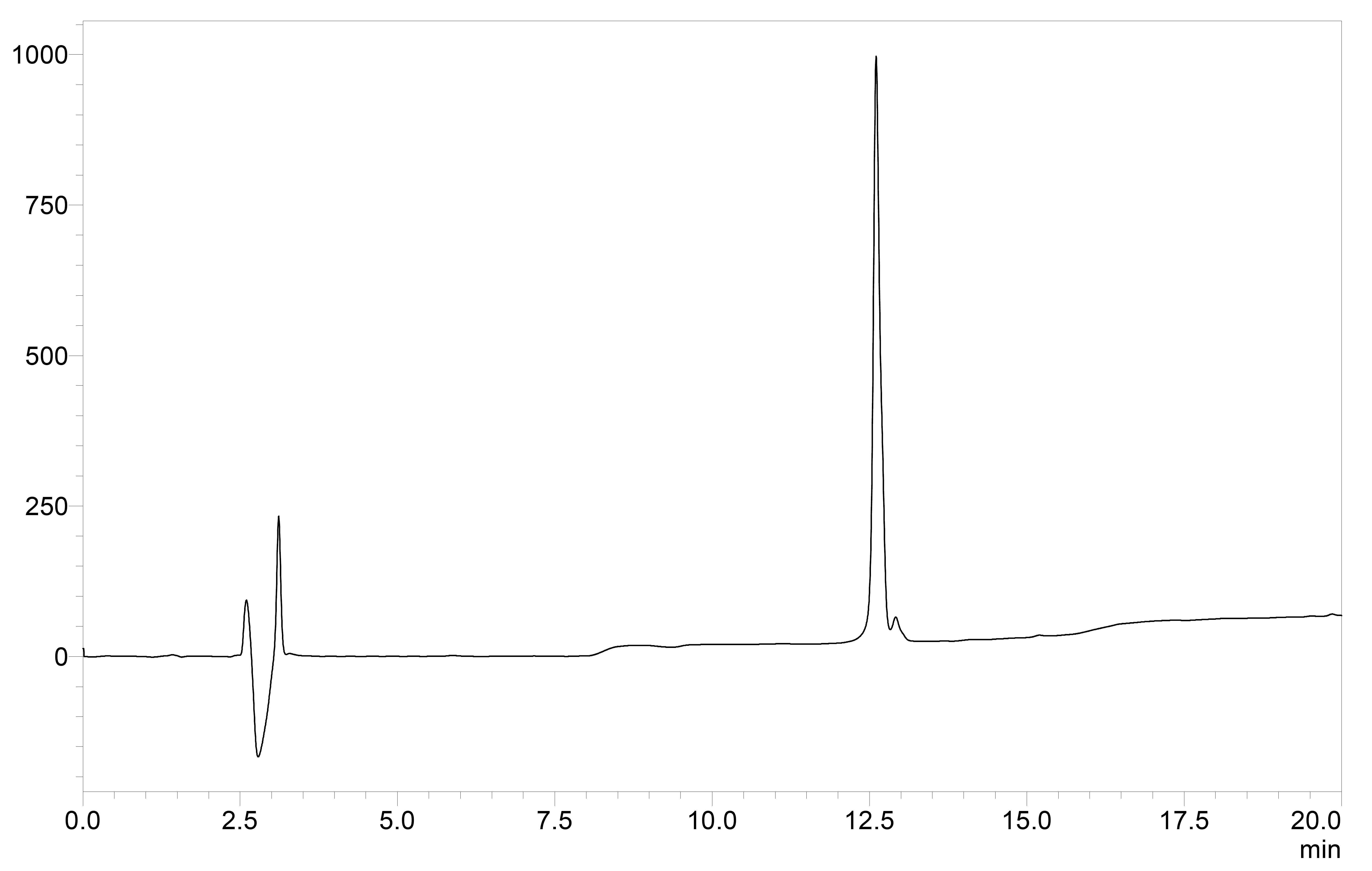

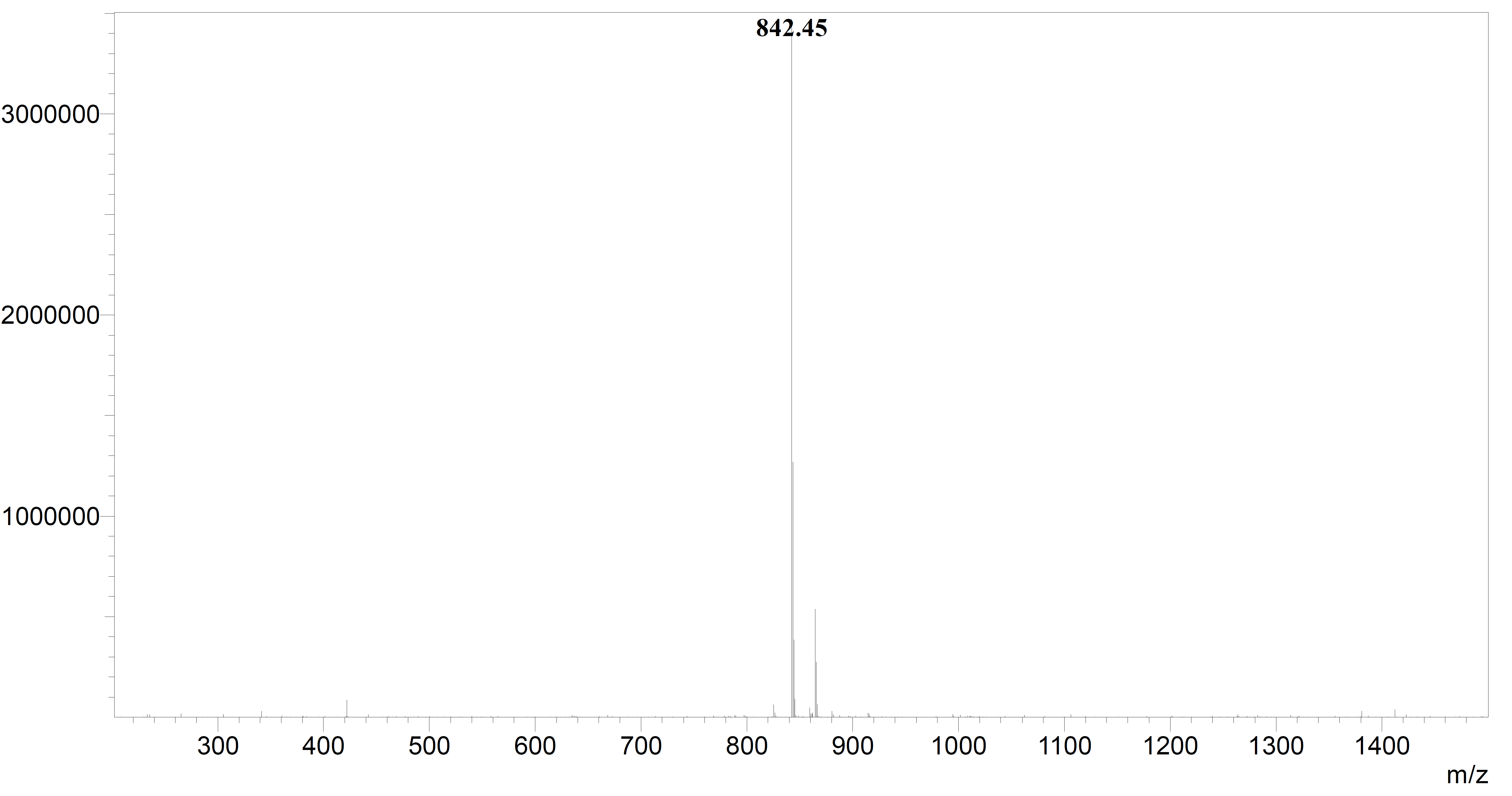

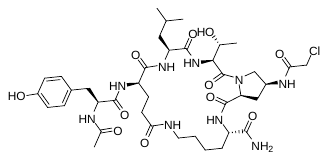
**APOD51** 32 mg, crude yield: 37%, purity: ≥ 95%, tR 12.78 min, (analytical HPLC, 10 to 90% acetonitrile (0.1% TFA) in water (0.1% TFA) over 20 min, flow rate of 1.0 mL/min); HRMS (ESI-MS): Calculated: 864.40 for C_39_H_59_ClN_9_O_11_ [M+H]^+^, found: 864.20.

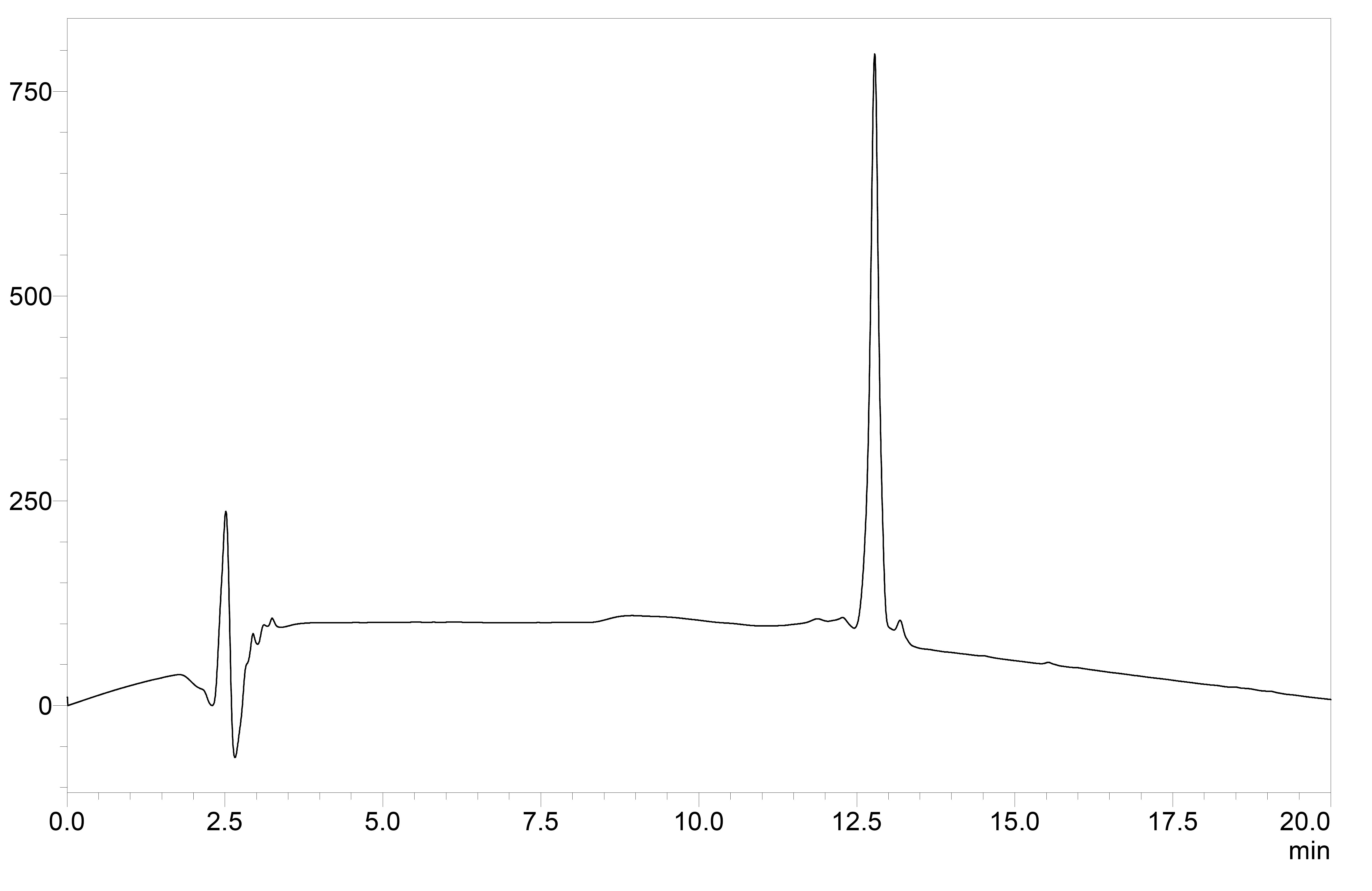

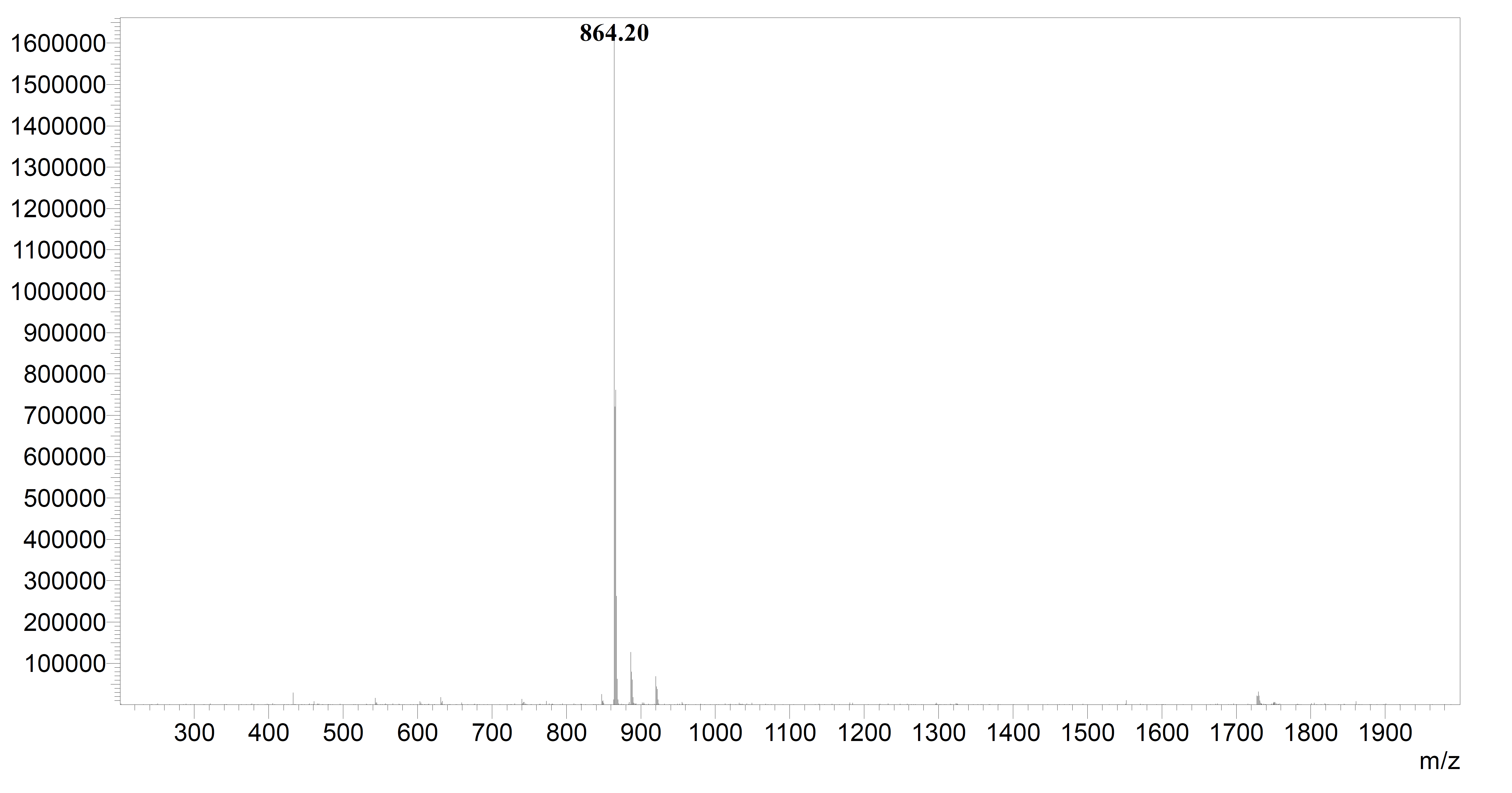

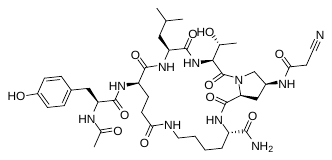
**APOD52** 38 mg, crude yield: 44 %, purity: ≥ 95%, tR 12.69 min, (analytical HPLC, 10 to 90% acetonitrile (0.1% TFA) in water (0.1% TFA) over 20 min, flow rate of 1.0 mL/min); ESI-MS: Calculated: 855.44 for C_40_H_59_N_10_O_11_ [M+H]^+^, found: 855.40.

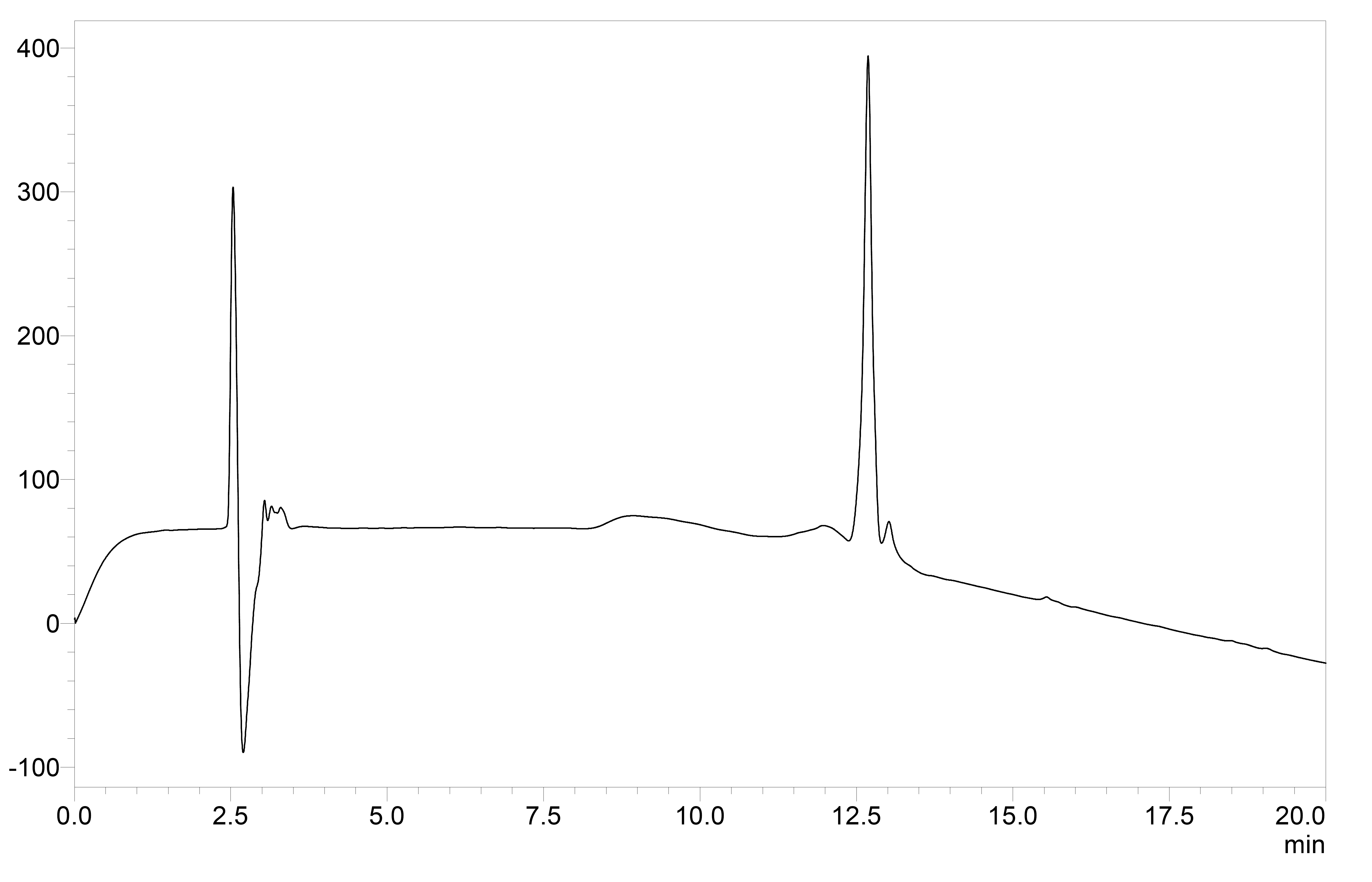

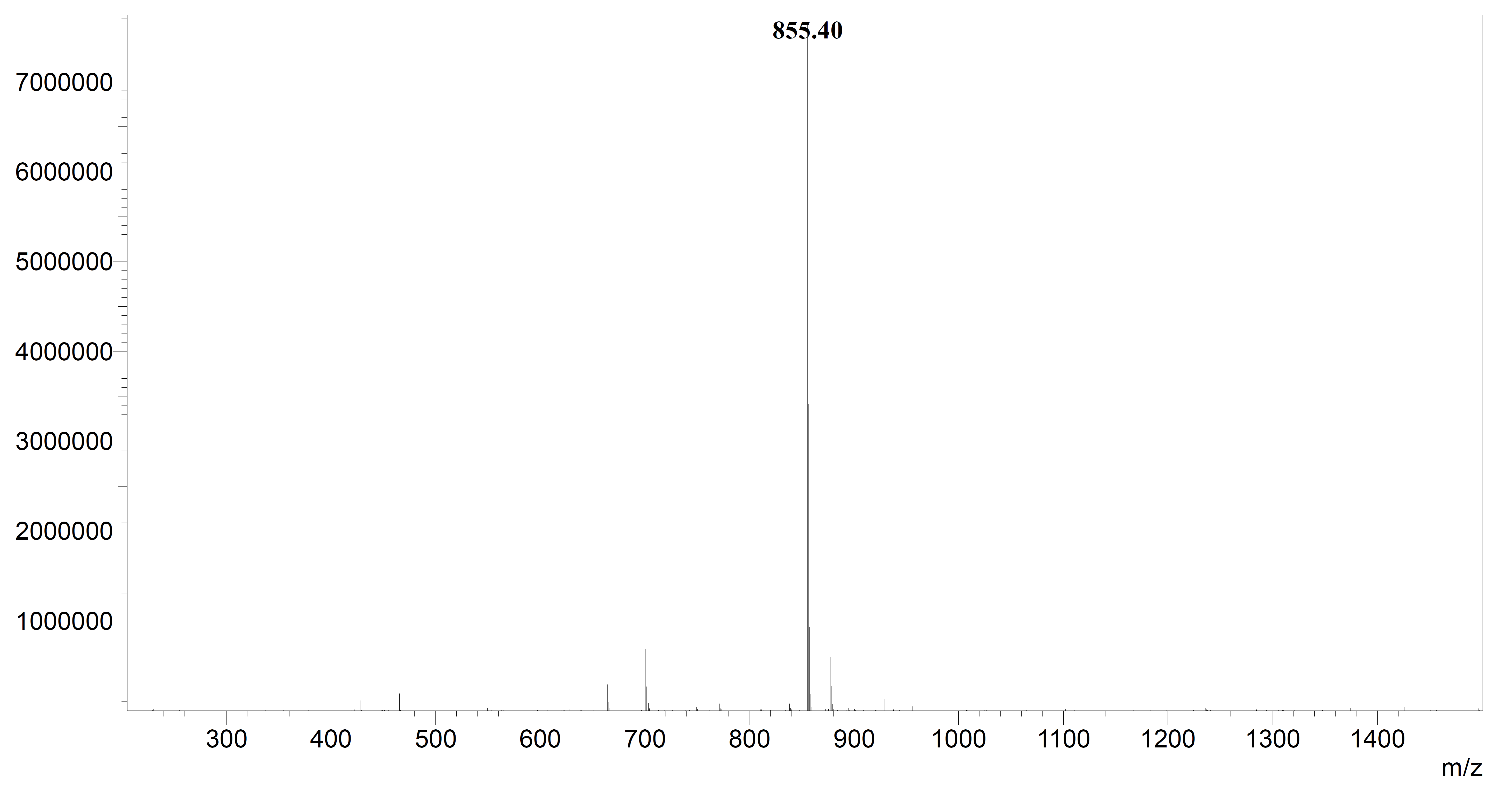

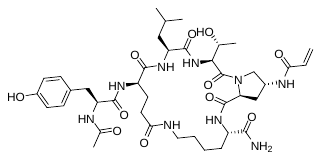
**APOD53** 40 mg, crude yield: 47 %, purity: ≥ 95%, tR 12.88 min, (analytical HPLC, 10 to 90% acetonitrile (0.1% TFA) in water (0.1% TFA) over 20 min, flow rate of 1.0 mL/min); HRMS (ESI-MS): Calculated: 842.96 for C_40_H_60_N_9_O_11_ [M+H]^+^, found: 842.25.

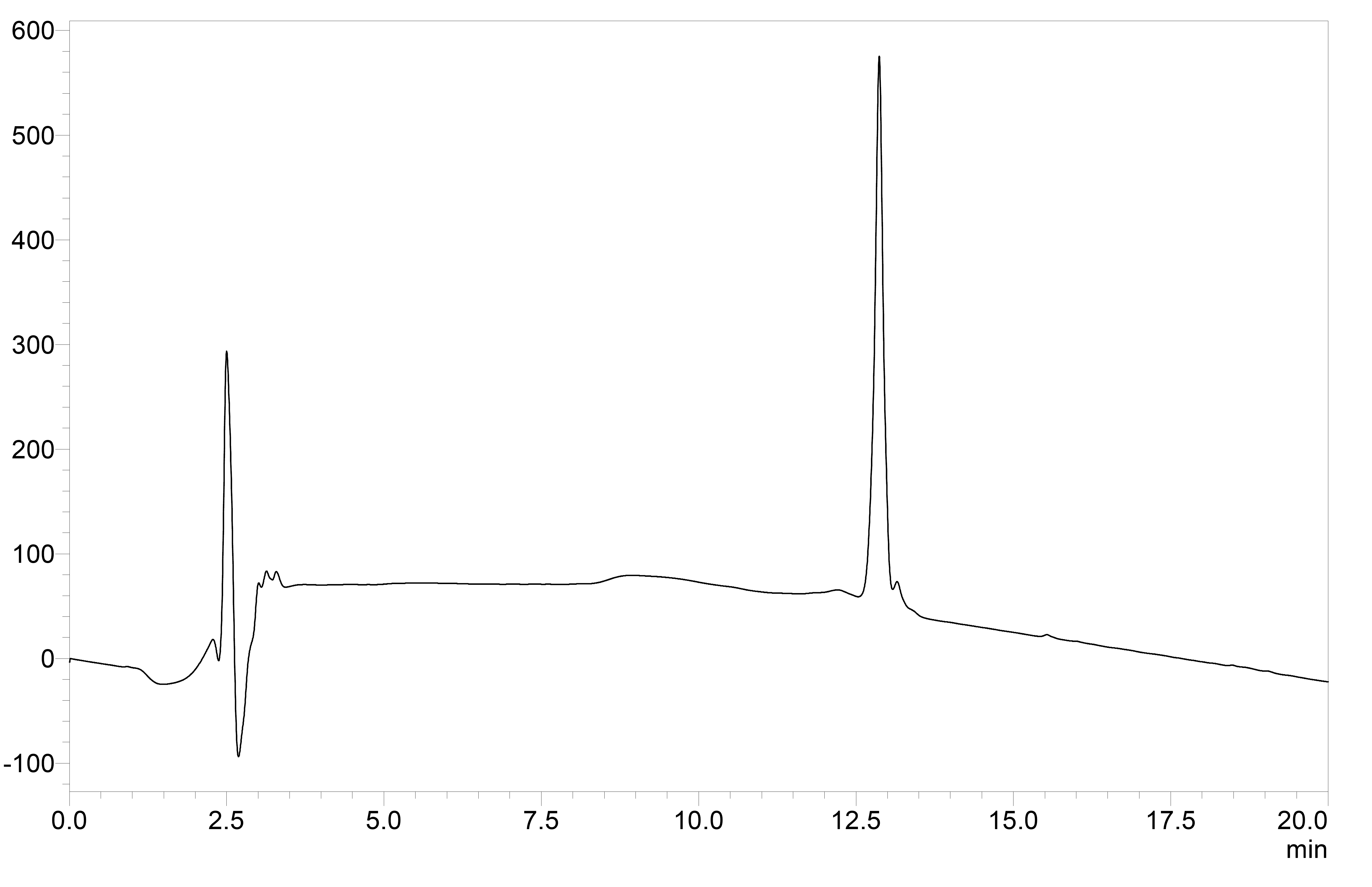

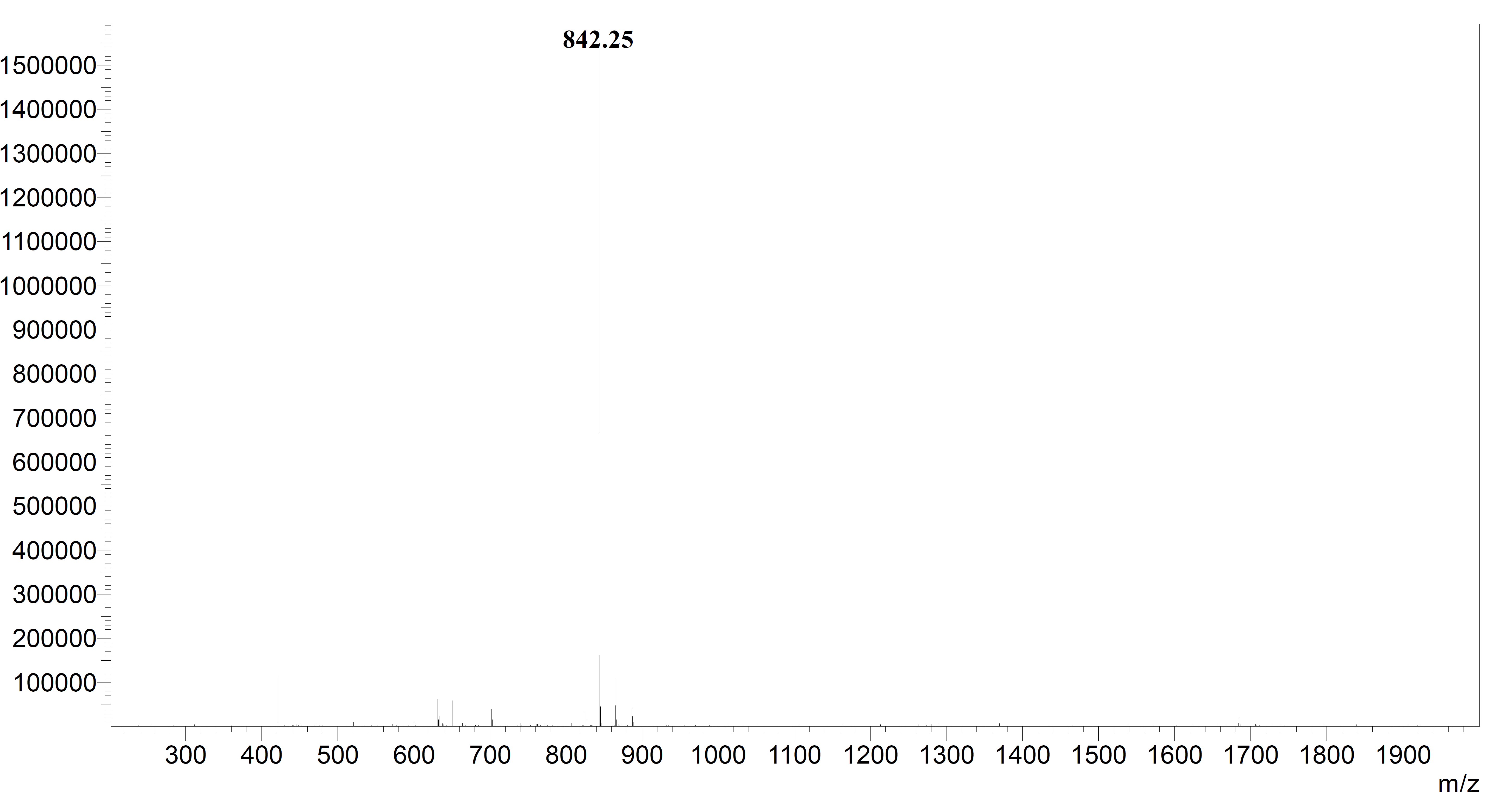

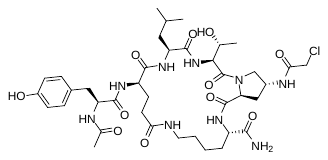
**APOD54** 33 mg, crude yield: 38 %, purity: ≥ 95%, tR 13.08 min, (analytical HPLC, 10 to 90% acetonitrile (0.1% TFA) in water (0.1% TFA) over 20 min, flow rate of 1.0 mL/min); HRMS (ESI-MS): Calculated: 864.40 for C_39_H_59_ClN_9_O_11_ [M+H]^+^, found: 864.25

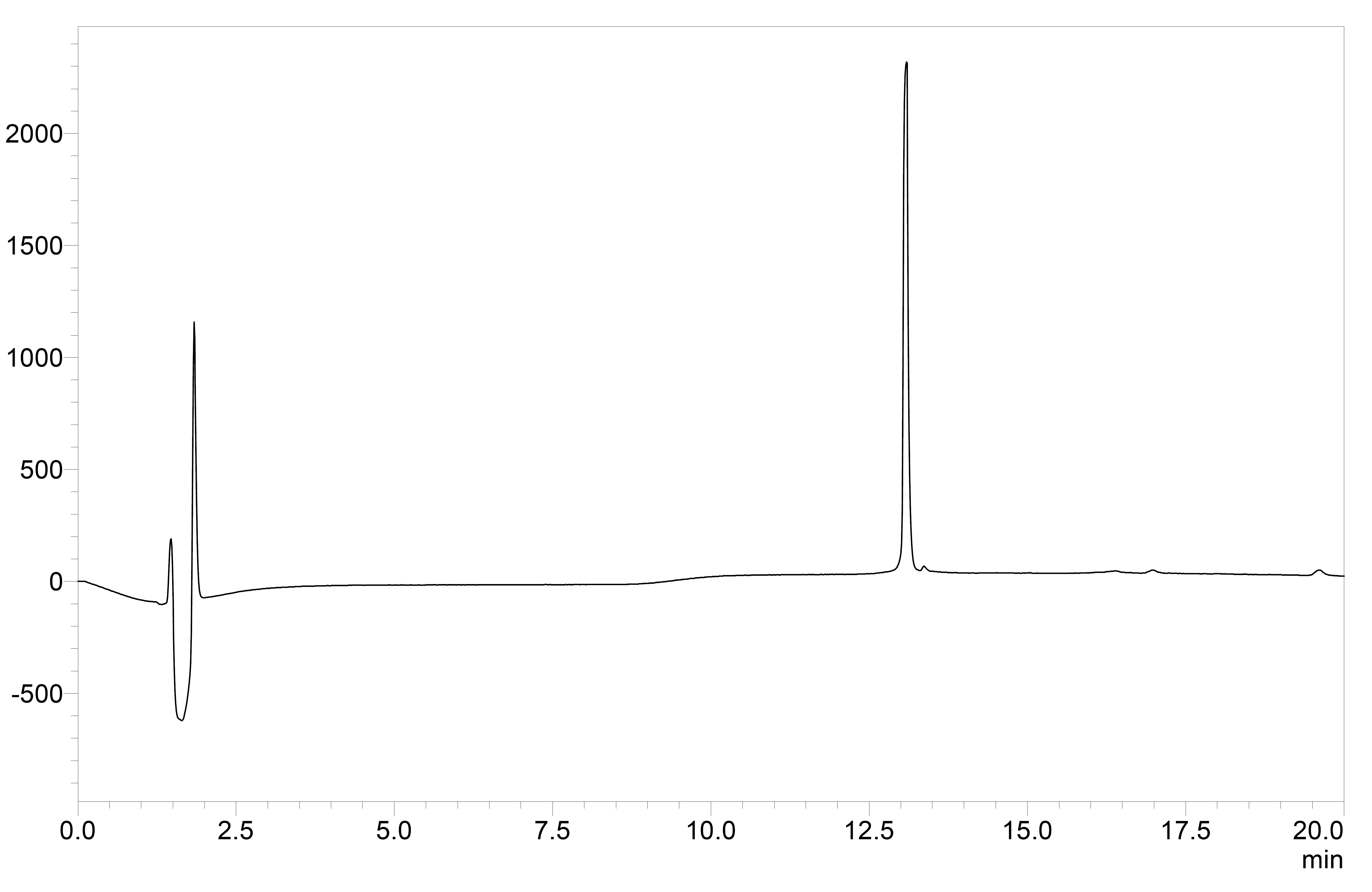

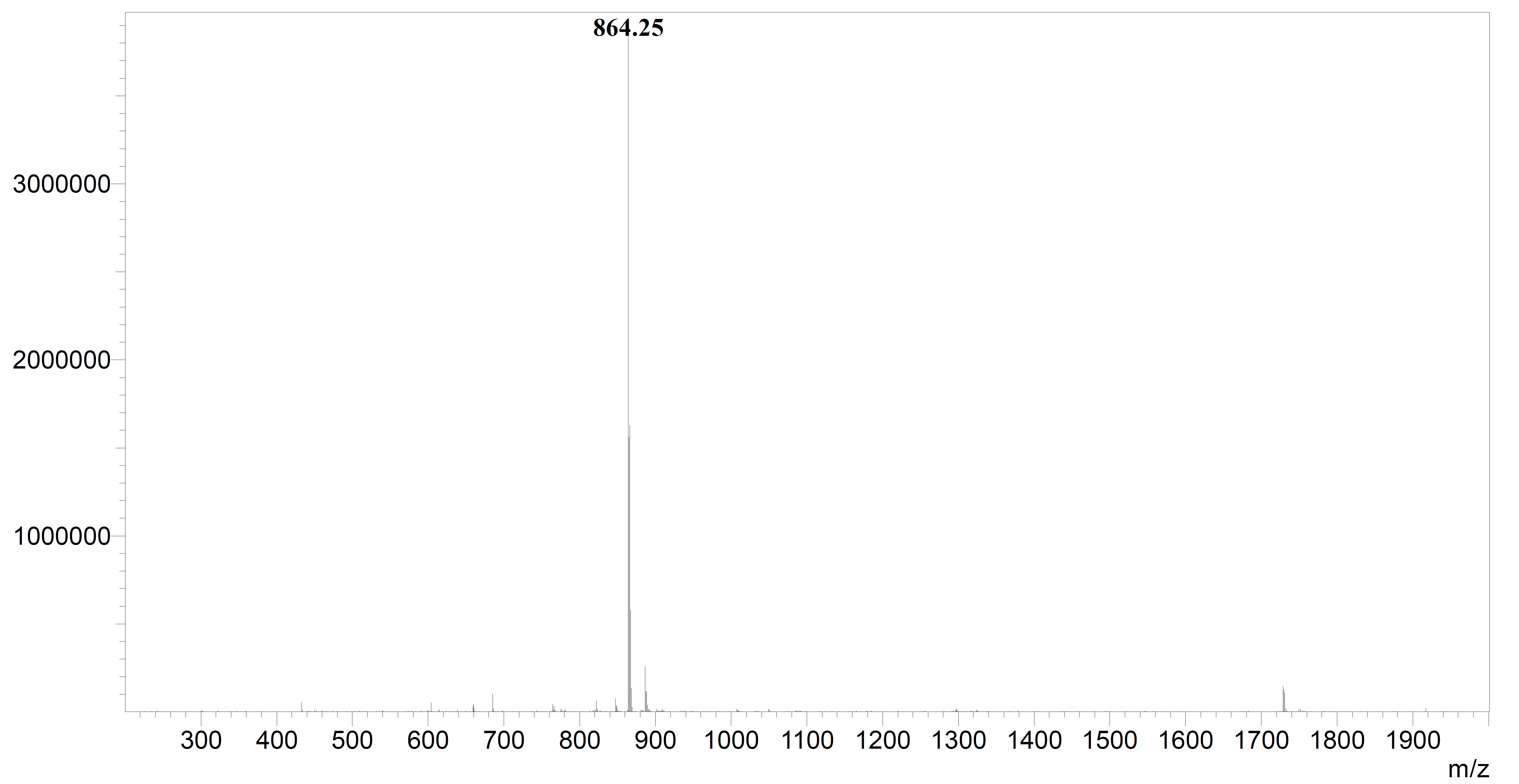

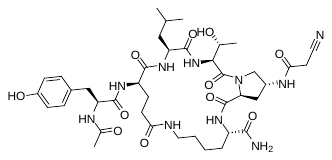
**APOD55** 38 mg, crude yield: 45 %, purity: ≥ 95%, tR 12.90 min, (analytical HPLC, 10 to 90% acetonitrile (0.1% TFA) in water (0.1% TFA) over 20 min, flow rate of 1.0 mL/min); HRMS (ESI-MS): Calculated: 855.44 for C_40_H_59_N_10_O_11_ [M+H]^+^, found: 855.25.

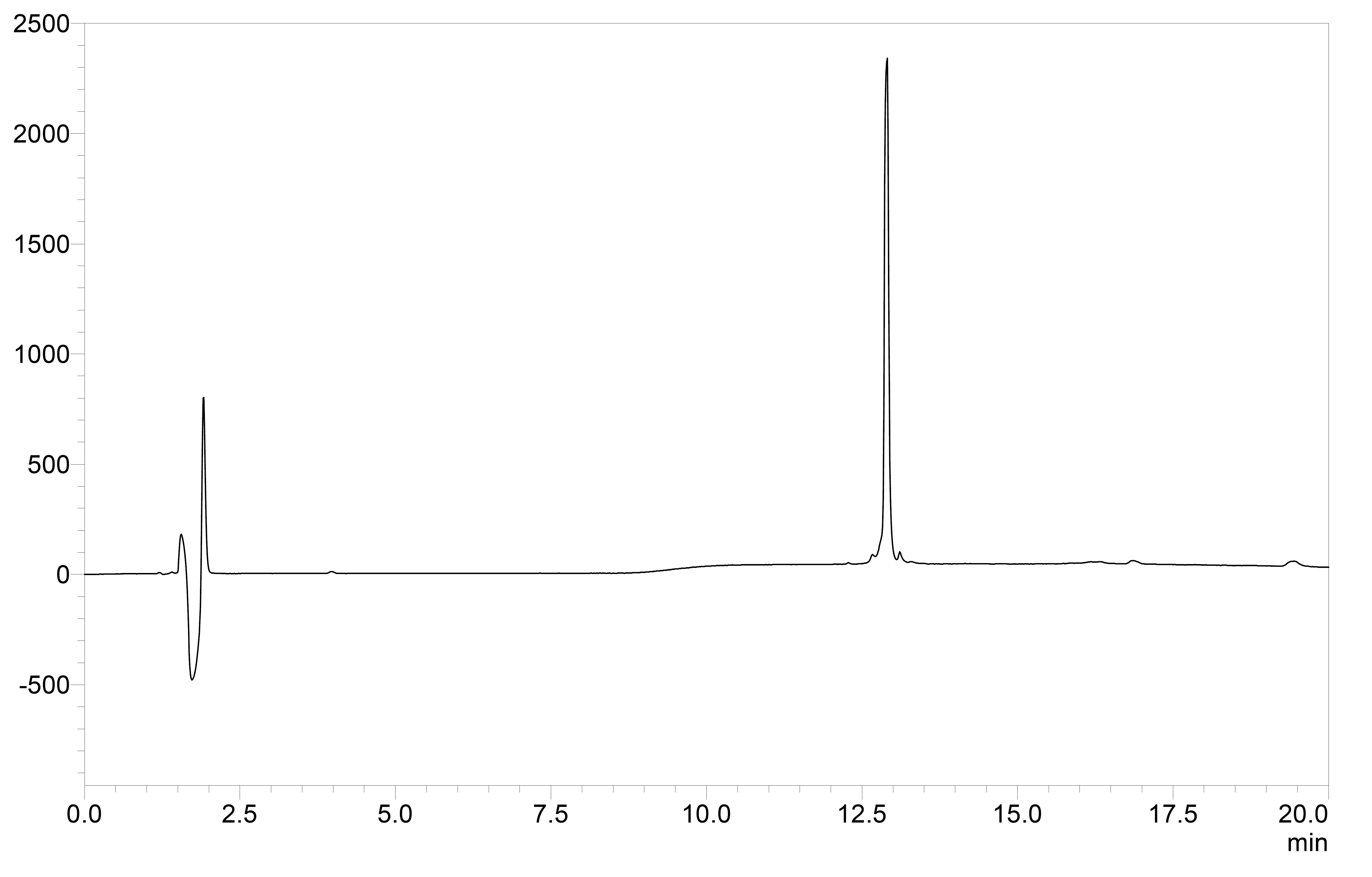

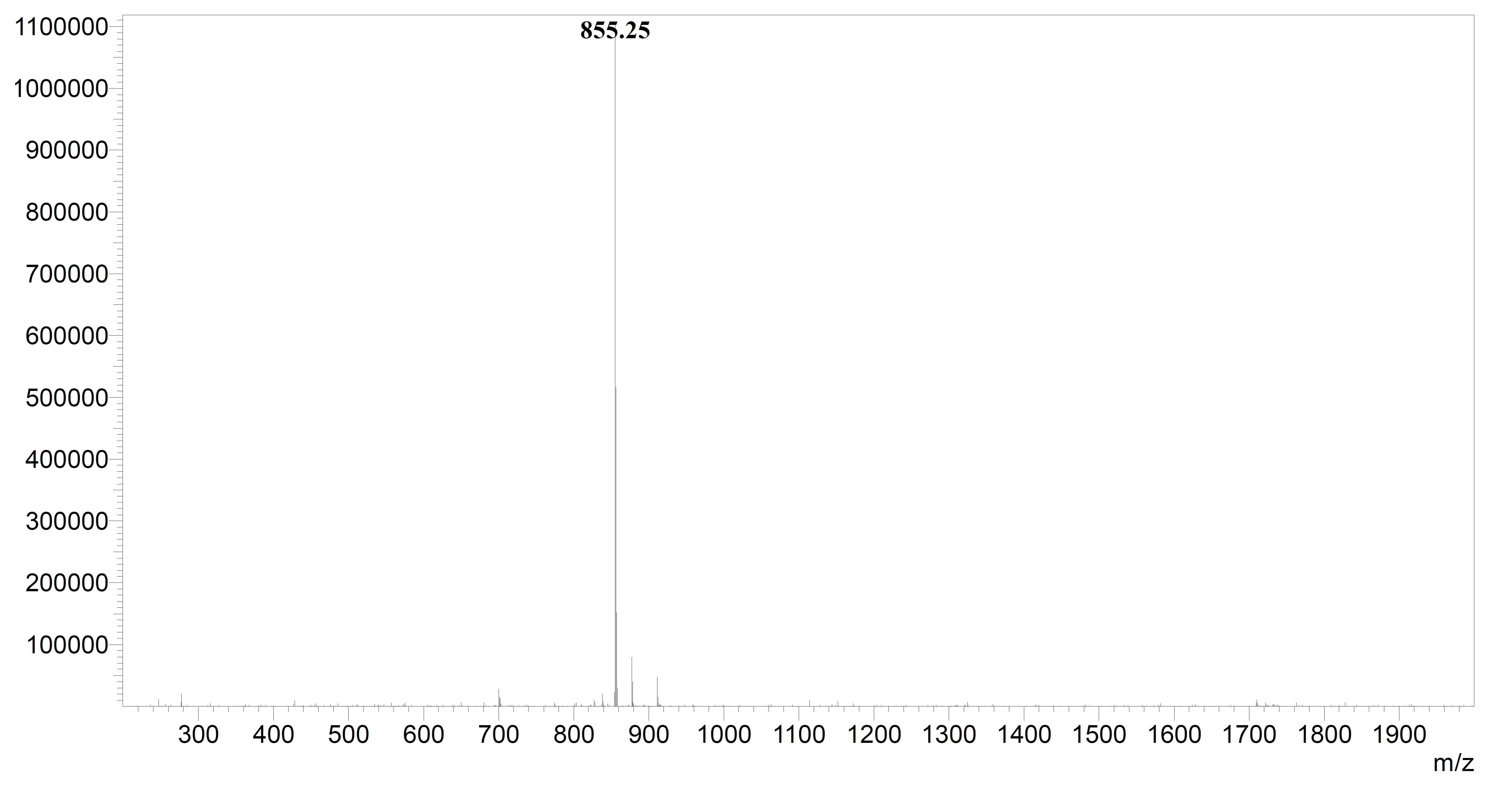

**sFigure 1.** HPLC chromatograms of RHPS4, STEREO 8, APOD41, and APOD50-55.

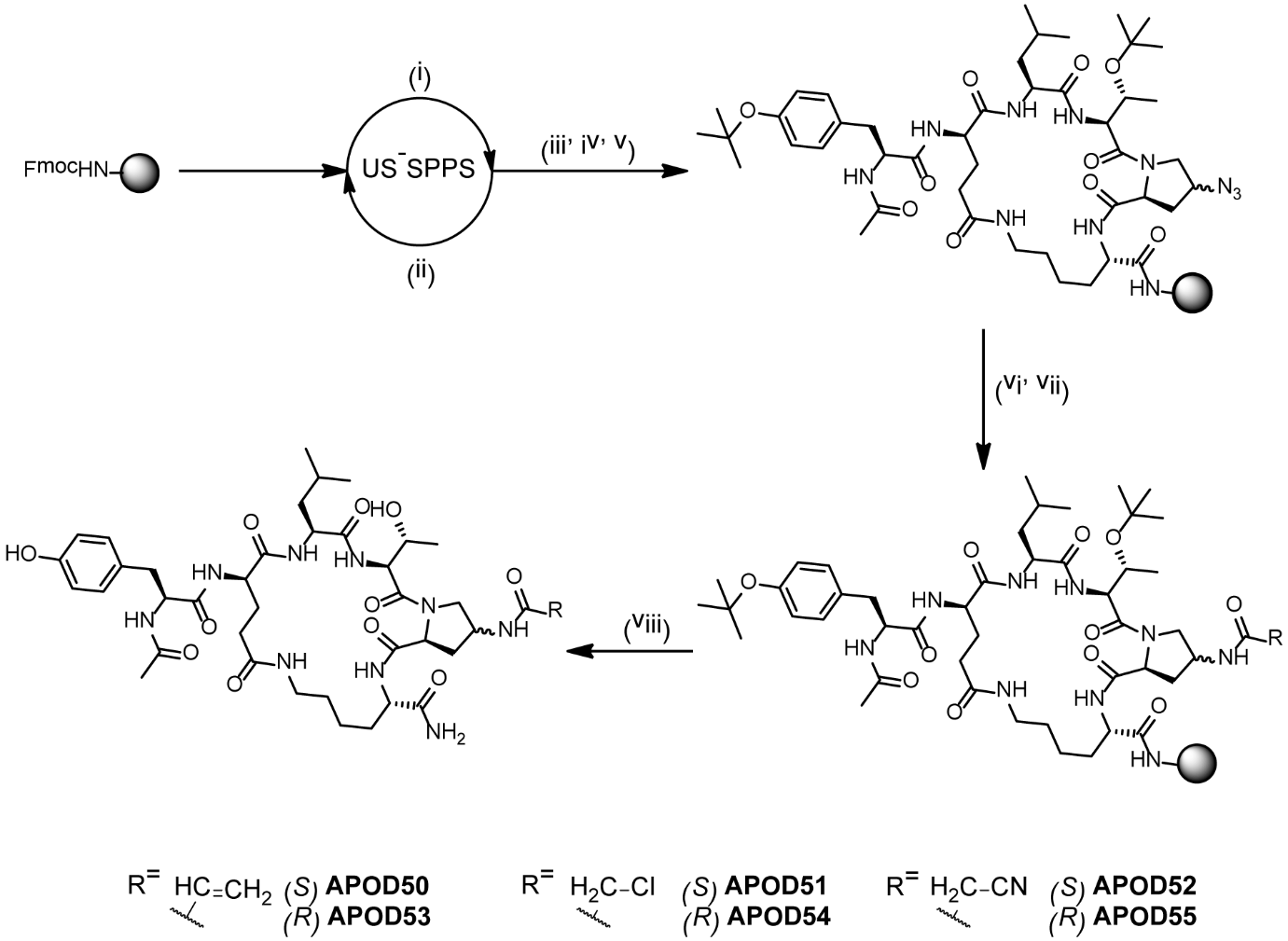

**sFigure 2.** General synthetic route for APOD50-APOD55. Reagents and conditions: i) piperidine 20% in DMF, 2 x 1 min, US irradiation; ii) Fmoc-AA-OH, COMU, OxymaPure, DIPEA, DMF, 5 min, US irradiation; iii) Ac2O, DIPEA, DMF, 10 min; iv) Pd(Ph3P)4, DMBA, DCM/DMF (1:1), 2 x 1h; v) PyAOP, HOAt, DIPEA, DMF, 16 h; vi) TCEP, THF/H2O (9:1), 12 h; vii) CH2CHCOCl, DIPEA, DMF, 30 min or ClCH2COCl, DIPEA, DMF, 30 min or CNCH2CO2H, COMU, OxymaPure, DIPEA, DMF; viii) TFA/TIS (95:5) 3 h.

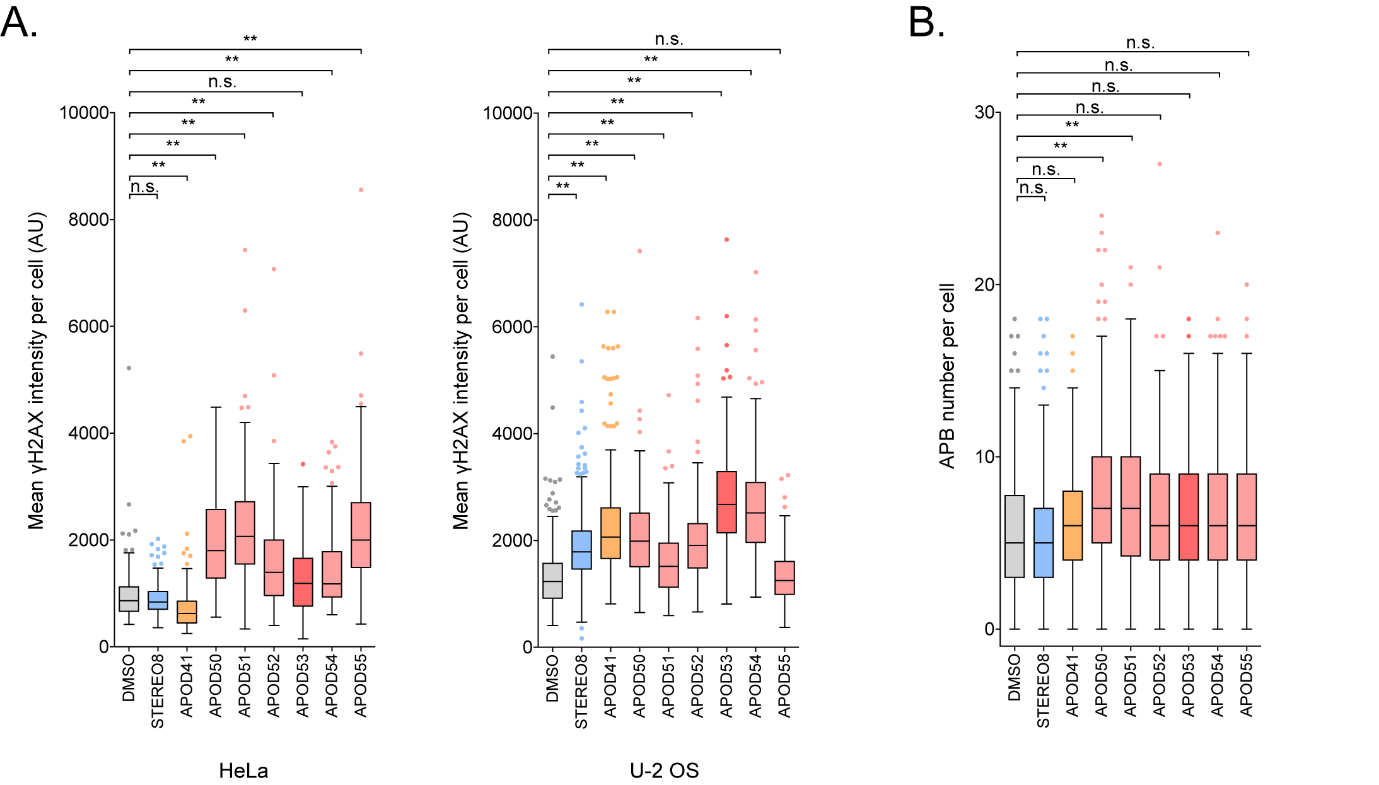
**sFigure 3.** Effects of APOD41 derivatives on genomic DNA damage and APB number. (A) Tukey box plots of mean γ-H2AX intensity per cell in HeLa and U-2 OS cells treated with 1 μM of APOD41 derivatives for 24 hrs. Out of three experiments, *n* = 150 cells scored in HeLa and *n* = 220 cells scored in U-2 OS per treatment, n.s. = non-significant, ***p* < 0.01, Kruskal-Wallis test. (B) Tukey box plots of APB frequency in U-2 OS cells treated with 1 μM of APOD41 derivatives for 24 hrs. Out of three experiments, *n* = 220 cells scored per treatment, n.s. = non-significant, **p* < 0.05, ***p* < 0.01, Kruskal-Wallis test.

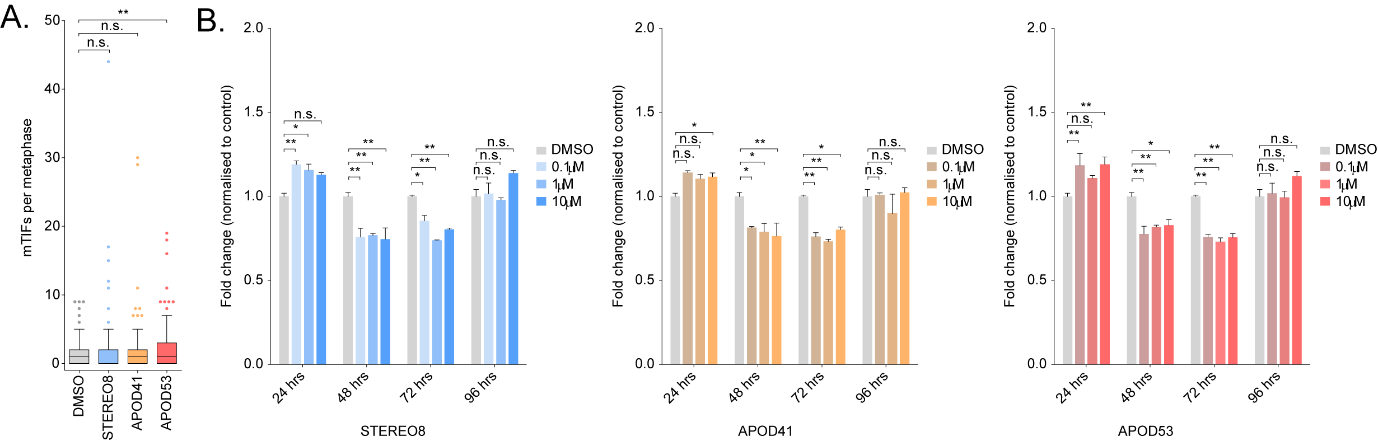

**sFigure 4.** Effects of the APOD41 derivative APOD53 on mortal cell line telomeric DNA damage and viability. (A) Tukey boxplots of metaphase-TIF in IMR-90 cells treated with 1 μM of APOD41 derivatives for 24 hrs. Out of three experiments, n ≥ 120 cells scored per treatment, n.s. = non-significant, **p < 0.005, Mann–Whitney test. (B) Quantification of WST-1 cell proliferation assays in IMR-90 with increasing dose over a 96 hr time period. Treated cells were normalized to the mean of DMSO control. Error bars represent the mean ± SEM from n = 3 experiments, n.s. = non-significant, *p < 0.05, **p < 0.01, ANOVA.

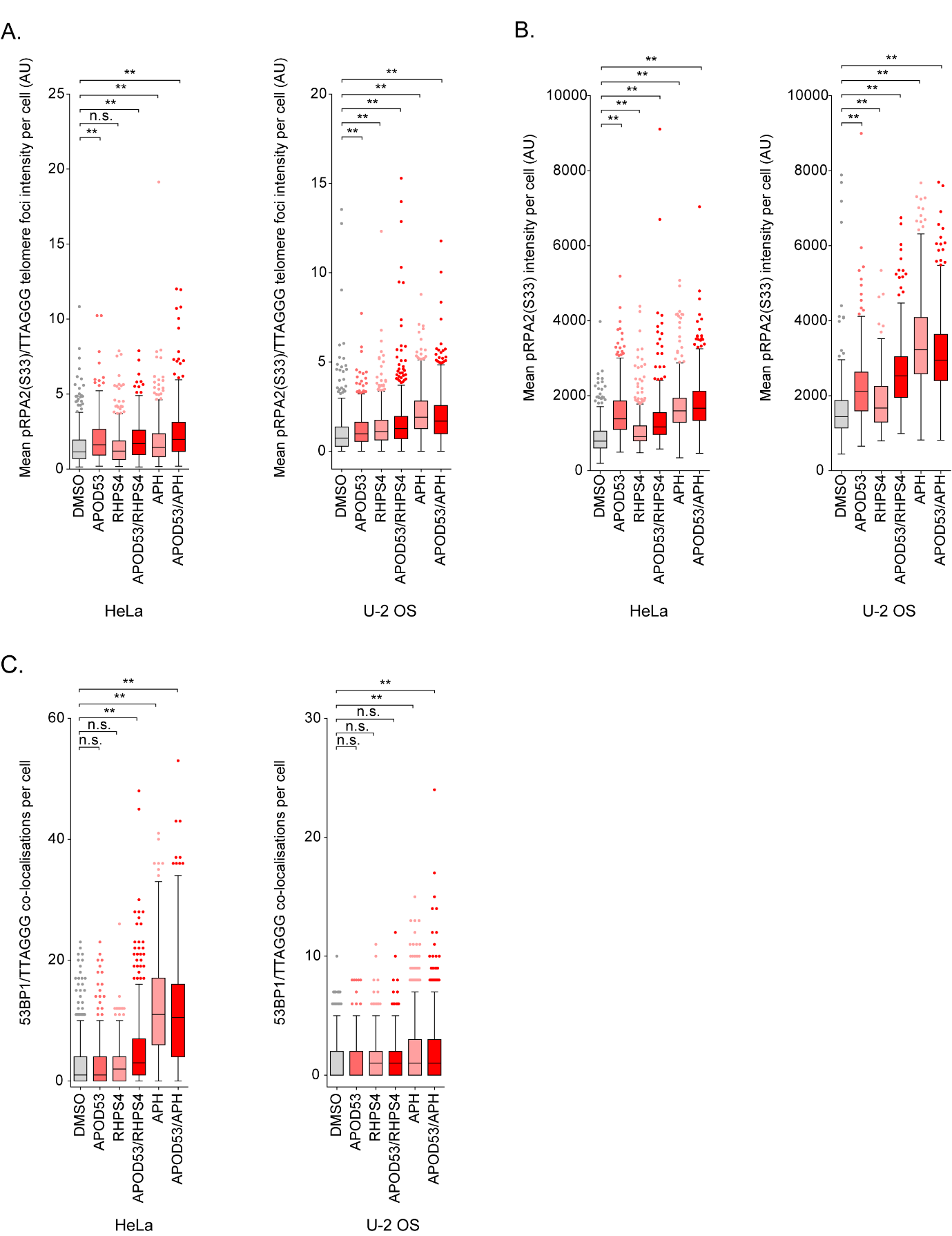

**sFigure 5.** Effects of APOD53 and RHPS4 co-treatment on pRPA2(S33) and 53BP1 signaling. (A) Tukey boxplots of pRPA2(S33)/TTAGGG telomere foci intensity per cell in HeLa and U-2 OS cells treated with 10 μM of APOD53 and 1 μM RHPS4 for 24 hrs. Out of three experiments, *n* = 550 cells scored in HeLa and *n* = 600 cells scored in U-2 OS per treatment, n.s. = non-significant, **p* < 0.05, ***p* < 0.01, Kruskal-Wallis test. (B) Tukey boxplots of mean pRPA2(S33) intensity per cell in HeLa and U-2 OS cells treated with 10 μM of APOD53 and 1 μM RHPS4 for 24 hrs. Out of three experiments, *n* = 550 cells scored in HeLa and *n* = 600 cells scored in U-2 OS per treatment, ***p* < 0.01, Kruskal-Wallis test. (C) Tukey boxplots of 53BP1/TTAGGG co-localizations in HeLa and U-2 OS cells treated with 10 μM of APOD53 and 1 μM RHPS4 for 24 hrs. Out of three experiments, *n* = 510 cells scored in HeLa and *n* = 570 cells scored in U-2 OS per treatment, n.s. = non-significant, ***p* < 0.01, Kruskal-Wallis test.

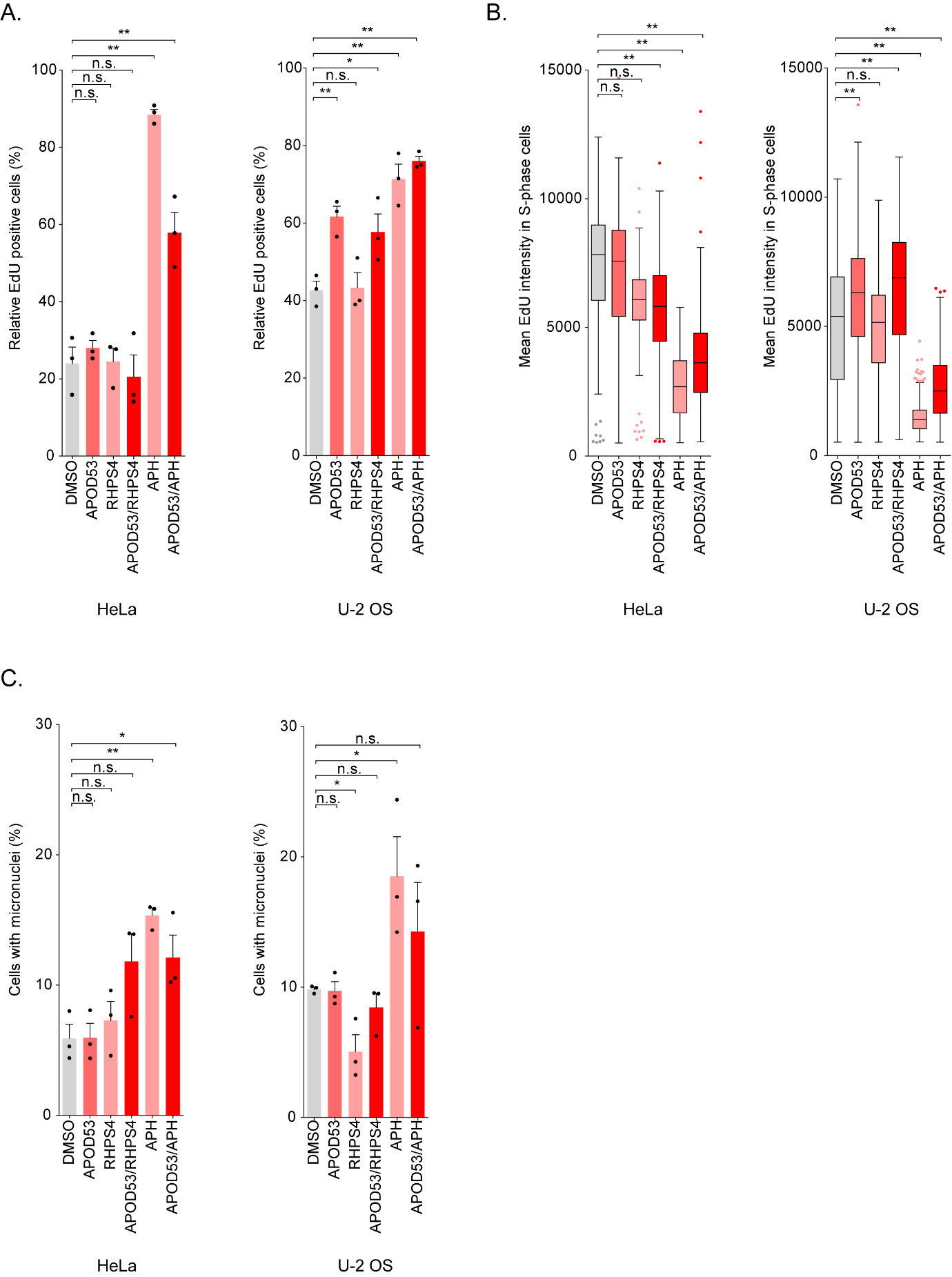

**sFigure 6.** Effects of APOD53 and RHPS4 co-treatment on genomic replication stress. (A) Mean number of EdU positive HeLa and U-2 OS cells treated with 10 μM of APOD53 and 1 μM RHPS4 for 24 hrs. Error bars represent the mean ± SEM from *n* = 3 experiments, *n* = 510 cells scored in HeLa and *n* = 550 cells scored in U-2 OS per treatment, n.s. = non-significant, **p* < 0.05, ***p* < 0.01, Student’s *t*-test. (B) Tukey boxplots of EdU intensity of EdU positive HeLa and U-2 OS S-phase cells treated with 10 μM of APOD53 and 1 μM RHPS4 for 24 hrs. Out of three experiments, *n* = 510 cells scored in HeLa and *n* = 550 cells scored in U-2 OS per treatment, n.s. = non-significant, ***p* < 0.01, Kruskal-Wallis test. (C) Mean number of cells with micronuclei in HeLa and U-2 OS cells treated with 10 μM of APOD53 and 1 μM RHPS4 for 24 hrs. Out of three experiments, *n* = 550 cells scored in HeLa and *n* = 600 cells scored in U-2 OS per treatment, n.s. = non-significant, **p* < 0.05, ***p* < 0.01, Student’s *t*-test.

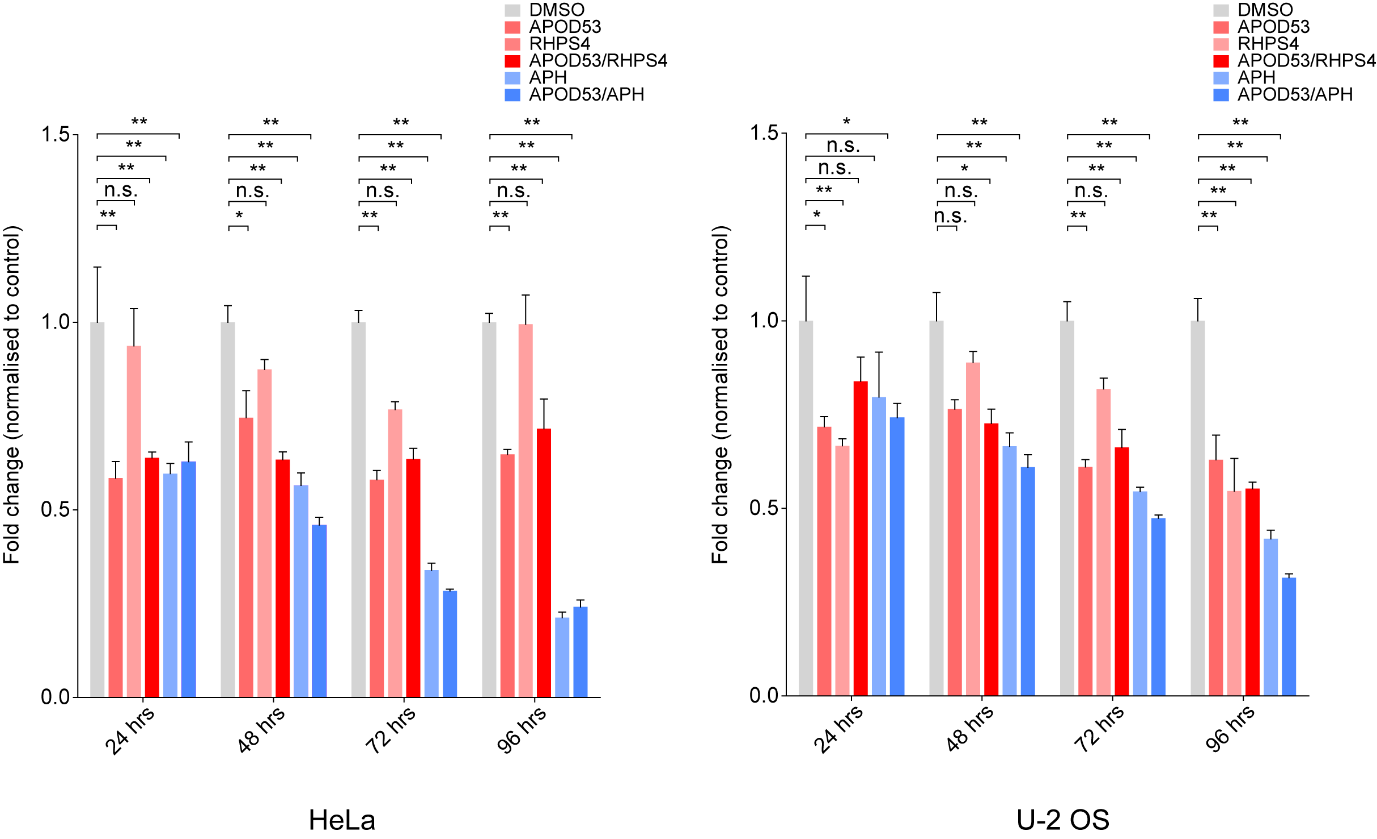

**sFigure 7.** Quantification of WST-1 cell proliferation assays in HeLa and U-2 OS treated with 10 μM of APOD53 and 1 μM RHPS4 over a 96 hr time period. Treated cells were normalized to the mean of DMSO control. Error bars represent the mean ± SD from *n* = 3 experiments.
